# Dynamic evolution of great ape Y chromosomes

**DOI:** 10.1101/2020.01.21.913863

**Authors:** Monika Cechova, Rahulsimham Vegesna, Marta Tomaszkiewicz, Robert S. Harris, Di Chen, Samarth Rangavittal, Paul Medvedev, Kateryna D. Makova

## Abstract

The mammalian male-specific Y chromosome plays a critical role in sex determination and male fertility. However, because of its repetitive and haploid nature, it is frequently absent from genome assemblies and remains enigmatic. The Y chromosomes of great apes represent a particular puzzle: their gene content is more similar between human and gorilla than between human and chimpanzee, even though human and chimpanzee shared a more recent common ancestor. To solve this puzzle, here we constructed a dataset including Ys from all extant great ape genera. We generated assemblies of bonobo and orangutan Ys, from short and long sequencing reads, and aligned them with the publicly available human, chimpanzee and gorilla Y assemblies. Analyzing this dataset, we found that the genus *Pan*, including chimpanzee and bonobo, experienced accelerated substitution rates. Additionally, *Pan* also exhibited elevated gene death rates. These observations are consistent with high levels of sperm competition in *Pan*. Furthermore, we inferred that the great ape common ancestor already possessed multi-copy sequences homologous to most human and chimpanzee palindromes. Nonetheless, each species also acquired distinct ampliconic sequences. We also detected increased chromatin contacts between and within palindromes (from Hi-C data), likely facilitating gene conversion and structural rearrangements. Moreover, our ENCODE data analysis suggested that Y palindromes exist to promote gene conversion preventing degradation of not only genes, as is commonly believed, but also gene regulatory sites. Our results highlight the dynamic mode of Y chromosome evolution, and open avenues for studies of male-specific dispersal in endangered great ape species.

## Introduction

The mammalian male-specific sex chromosome—the Y—is vital for sex determination and male fertility, and is a useful marker for population genetics studies. It carries *SRY* encoding the testis-determining factor that initiates male sex determination (1). The human Y also harbors azoospermia factor regions, deletions of which can cause infertility (2). Y chromosome sequences have been used to analyze male dispersal (3) and hybridization sex bias (4) in natural populations. Thus, the Y is important biologically and its sequences have critical practical implications. Moreover, study of the Y is needed to obtain a complete picture of mammalian genome evolution. Yet, due to its repetitive and haploid nature, the Y has been assembled for only a handful of mammalian species (5).

Among great apes, the Y has so far been assembled only in human (6), chimpanzee (7), and gorilla (8). A comparative study of these Y assemblies (8) uncovered some unexpected patterns which could not be explained with the data from three species alone. Despite a recent divergence of these species (~7 million years ago, MYA (9)), their Y chromosomes differ enormously in size and gene content, in sharp contrast with the stability of the rest of the genome. For instance, the chimpanzee Y is only half the size of the human Y, and the percentage of gene families shared by these two chromosomes (68%) that split ~6 MYA (9) is similar to that shared by human and chicken autosomes that split ~310 MYA (7). Puzzlingly, in terms of shared genes and overall architecture, the human Y is more similar to the gorilla Y than to the chimpanzee Y even though human and chimpanzee have a more recent common ancestor (8). Y chromosomes from additional great ape species should be sequenced to understand whether high interspecific variability in gene content and architecture is characteristic of all great ape Ys.

All great ape Y chromosomes studied thus far include *pseudoautosomal regions (PARs)*, which recombine with the X chromosome, and *male-specific* X-degenerate, ampliconic, and heterochromatic regions, which evolve as a single linkage group (6–8). The *X-degenerate regions* are composed of segments with different levels of homology to the X chromosome—*strata*, corresponding to stepwise losses of recombination on the Y. Because of lack of recombination, X-degenerate regions are expected to accumulate gene-disrupting mutations, however this has not been examined in detail. The *ampliconic regions* consist of repetitive sequences that have >50% identity to each other and contain *palindromes—*inverted repeats (separated by a spacer) up to several megabases long whose arms are >99.9% identical (6). Palindromes are thought to evolve to allow intrachromosomal (Y-Y) gene conversion (10) which rescues the otherwise non-recombining male-specific regions from deleterious mutations (11). We presently lack knowledge about how conserved palindrome sequences are across great apes. The *heterochromatic regions* are rich in satellite repeats (12). In general, X-degenerate regions are more conserved, whereas ampliconic regions are prone to rearrangements, and heterochromatic regions evolve very rapidly, among species (7, 8, 12). However, the evolution of great ape Y chromosomes outside of human, chimpanzee, and gorilla has not been explored.

Known Y chromosome protein-coding genes are located in either the X-degenerate or ampliconic regions. *X-degenerate genes* (16 on the human Y) are single-copy, ubiquitously expressed genes with housekeeping functions (13). Multi-copy *ampliconic genes* (nine gene families on the human Y, eight of which—all but *TSPY*—are located in palindromes) are expressed only in testis and function during spermatogenesis (6). Some human Y genes are deleted or pseudogenized in other great apes, and thus are not essential for all species (8, 14). To illuminate genes essential for male reproduction in non-human great apes, all of which are endangered species, a cross-species analysis of Y gene content evolution is needed.

Here we compared the Y chromosomes in five species representing all four great ape genera: the human *(Homo)* lineage diverged from the chimpanzee (*Pan*), gorilla *(Gorilla)*, and orangutan (*Pongo*) lineages ~6 MYA, ~7 MYA, and ~13 MYA, respectively (9), and the bonobo and chimpanzee lineages (belonging to the genus *Pan)*, which diverged ~1 MYA (15). We produced draft assemblies of the bonobo and Sumatran orangutan Ys, and combined them with the human, chimpanzee and gorilla Y assemblies (6–8) to construct great ape Y multi-species alignments. This comprehensive data set enabled us to answer several pivotal questions about evolution of great ape Y chromosomes. First, we assessed lineage-specific substitution rates, and identified species experiencing significant rate acceleration. Second, we evaluated the conservation of palindromic sequences, and examined chromatin interactions within ampliconic regions. Third, we determined interspecific gene content turnover. Our results highlight the highly dynamic nature of great ape Y chromosome evolution.

## Results

### Assemblies

To obtain Y chromosome assemblies for all major great ape lineages, we augmented publicly available human and chimpanzee assemblies (6, 7) by producing draft bonobo and Sumatran orangutan (henceforth called ‘orangutan’) assemblies, and by improving the gorilla assembly (8), of Y male-specific regions (**Fig. S1**, see Methods for details). The resulting assemblies (henceforth called ‘Y assemblies’) were of high quality, as evidenced by their high degree of homology to the human and chimpanzee Ys (**Fig. S2**) and by the presence of sequences of most expected (14) homologs of human Y genes (**Fig. S3**). They also were of sufficient continuity (**Table S1**), particularly taking the highly repetitive structure of the Y into account.

### Ampliconic and X-degenerate scaffolds

To determine which scaffolds are ampliconic and which are X-degenerate in our bonobo, gorilla, and orangutan Y assemblies (such annotations are already available for the human and chimpanzee Ys (6, 7)), we developed a classifier which combines the copy count in the assemblies with mapping read depth information from whole-genome sequencing of male individuals (**Note S1**). This approach was needed as ampliconic regions can be collapsed in assemblies based on next-generation sequencing data (5). Using this classifier, we identified 12.5 Mb, 10.0 Mb, and 14.5 Mb of X-degenerate scaffolds in bonobo, gorilla, and orangutan, respectively. The length of ampliconic regions was more variable: 10.8 Mb in bonobo, 4.0 Mb in gorilla, and 2.2 M in orangutan. Due to potential collapse of repeats, we might have underestimated the true lengths of ampliconic regions. However, their length estimates are expected to reflect their complexity: e.g., the complexity might be low in the orangutan Y, which is consistent with a high read depth in its Y ampliconic scaffolds (**Fig. S4**) and with its longgene-harboring repetitive arrays found previously cytogenetically (16).

### Alignments

We aligned the sequences of the Y chromosomes from five great ape species (see Methods for details). The resulting multi-species alignment allowed us to identify species-specific sequences, sequences shared by all species, and sequences shared by some but not all species (**Fig. S5, Table S2A**). These results were confirmed by pairwise alignments (**Table S2B**). For instance, as was shown previously (8), the gorilla Y had the highest percentage of its sequence aligning to the human Y (75.7% and 89.6% from multi-species and pairwise alignments, respectively). In terms of sequence identity (**Tables S2C-D**), the chimpanzee and bonobo Ys were most similar to each other (99.2% and ~98% from multi-species and pairwise alignments, respectively), while the orangutan Y had the lowest identity to any other great ape Y chromosomes (~93-94% and ~92% from multi-species and pairwise alignments, respectively). From multi-species alignments (**Table S2C**), the human Y was most similar in sequence to the chimpanzee or bonobo Ys (97.9% and 97.8%, respectively), less similar to the gorilla Y (97.2%), and the least similar to the orangutan Y (93.6%), in agreement with the accepted phylogeny of these species (9). The pairwise alignments confirmed this trend (**Table S2D**). These results argue against incomplete lineage sorting at the male-specific Y chromosome locus in great apes.

### Substitution rates on the Y

We next asked whether the chimpanzee Y chromosome, whose architecture and gene content differ drastically from the human and gorilla Ys (8), experienced an elevated substitution rate. Using our multi-species Y chromosome alignment, we estimated substitution rates along the branches of the great ape phylogenetic tree (**Fig. 1A**; see Methods for details). A similar analysis was performed using an alignment of autosomes (**Fig. 1B**). A higher substitution rate on the Y than on the autosomes, i.e. male mutation bias (17), was found for each branch of the phylogeny (**Fig. 1**, **Note S2**). Notably, the Y-to-autosomal substitution rate ratio was higher in the *Pan* lineage, including the chimpanzee (1.76) and bonobo (1.76) lineages and the lineage of their common ancestor (1.91), than in the human lineage (1.48). These trends did not change after correcting for ancestral polymorphism (**Note S2**). We subsequently used a test akin to the relative rate test (18) and addressed whether the *Pan* lineage experienced more substitutions than the human lineage (**Table S3**). Using gorilla as an outgroup, we observed a significantly higher number of substitutions that occurred between chimpanzee and gorilla than between human and gorilla. The ratio of these two numbers was 1.006 (significantly different from 1, *p*<1×10^-5^, χ^2^-test) for autosomes, but was as high as 1.090 (*p*<1×10^-5^, χ^2^-test) for the Y. Similarly, we observed a higher number of substitutions that occurred between bonobo and gorilla than between human and gorilla. The ratio of these two numbers was 1.029 (*p*<1×10^-5^, χ^2^-test) for autosomes, but was as high as 1.114 (*p* <1×10^-5^, χ^2^-test) for the Y. In both cases, these ratios were significantly higher for the Y than for the autosomes (*p*<1×10^-5^ in both cases, χ^2^-test on contingency table). Thus, while the *Pan* lineage experienced an elevated substitution rate at both autosomes and the Y, this elevation was particularly strong on the Y.

**Figure 1.**
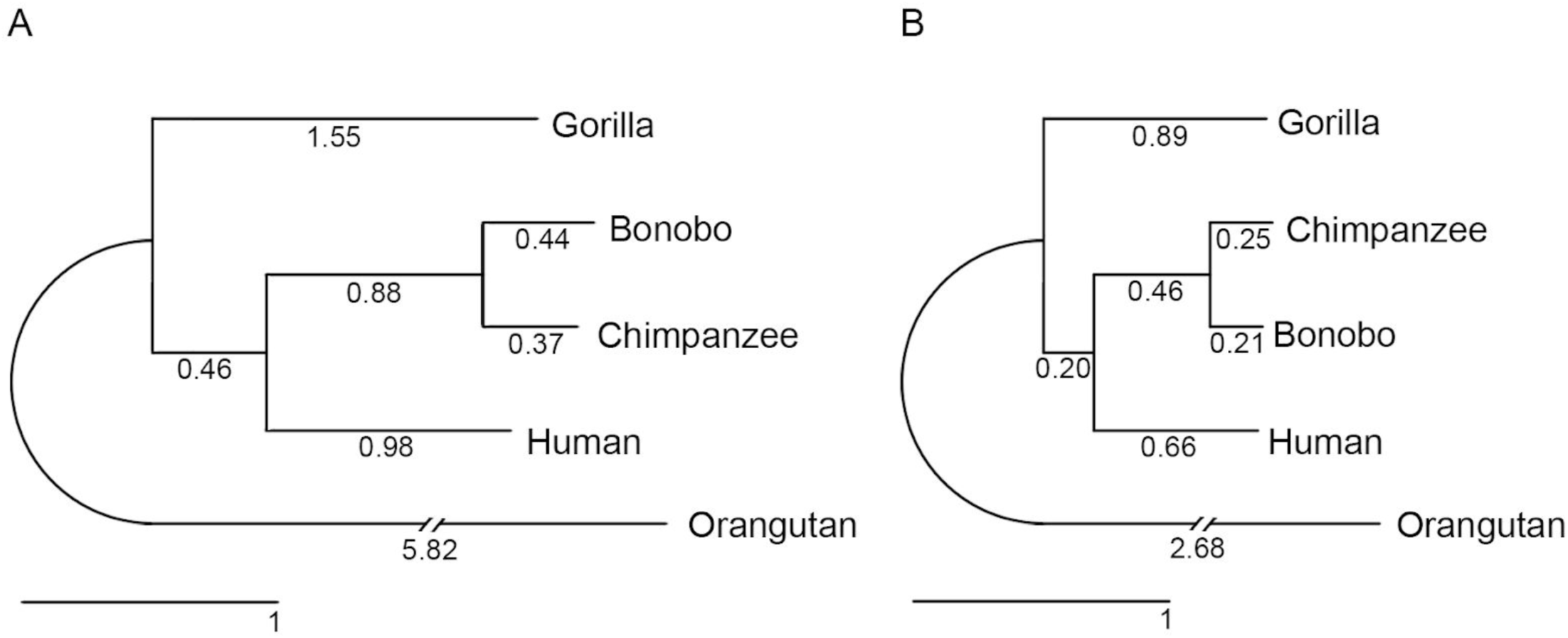
Phylogenetic tree of nucleotide sequences. (**A**) Y chromosome, and (**B**) Autosomes. Branch lengths (substitutions per 100 sites) were estimated from multi-species alignment blocks including all five species.

### Gene content evolution

Utilizing sequence assemblies and testis expression data (19), we evaluated gene content and the rates of gene birth and death on the Y chromosomes of five great ape species. First, we examined the presence/absence of homologs of human Y chromosome genes (16 X-degenerate genes + 9 ampliconic gene families = 25 gene families; for multi-copy ampliconic gene families, we were not studying copy number variation, but only presence/absence of a family in a species; **Fig. S6**). Such data were previously available for the chimpanzee Y, in which seven out of 25 human Y gene families became pseudogenized or deleted (7), and for the gorilla Y, in which only one gene family *(VCY)* out of 25 is absent (8). Here, we compiled the data for bonobo and orangutan. From the 25 gene families present on the human Y, the bonobo Y lacked seven *(HSFY, PRY, TBL1Y, TXLNGY, USP9Y, VCY*, and *XKRY)* and the orangutan Y lacked five *(TXLNGY, CYorf15A, PRKY, USP9Y*, and *VCY).* Second, our gene annotation pipeline did not identify novel genes in the bonobo and orangutan Y assemblies (**Note S3**), similar to previous results for the chimpanzee (7) and gorilla (8) Ys. Thus, we obtained the complete information about gene family content on the Y chromosome in five great ape species.

Using this information and utilizing the macaque Y chromosome (20) as an outgroup, we reconstructed gene content at ancestral nodes and studied the rates of gene birth and death (21) across the great ape phylogeny. Because X-degenerate and ampliconic genes might exhibit different trends, we analyzed them separately (**Fig. 2, Table S4**). Considering gene births, none were observed for X-degenerate genes, and only one (*VCY*, in the human-chimpanzee-bonobo common ancestor) was observed for ampliconic genes, leading to overall low gene birth rates. Considering gene deaths, three ampliconic gene families and three X-generate genes were lost by the chimpanzee-bonobo common ancestor, leading to death rates of 0.095 and 0.049 events/MY, respectively. Bonobo lost an additional ampliconic gene, whereas chimpanzee lost an additional X-degenerate gene, leading to death rates of 0.182 and 0.080 events/MY, respectively. In contrast, no deaths of either ampliconic or X-degenerate genes were observed in human and gorilla. Orangutan did not experience any deaths of X-degenerate genes, but lost four ampliconic genes. Its ampliconic gene death rate (0.021 events/MY) was still lower than that in the bonobo or in the bonobo-chimpanzee common ancestor. To summarize, the *Pan* genus exhibited the highest death rates for both X-degenerate and ampliconic genes across great apes.

**Figure 2.**
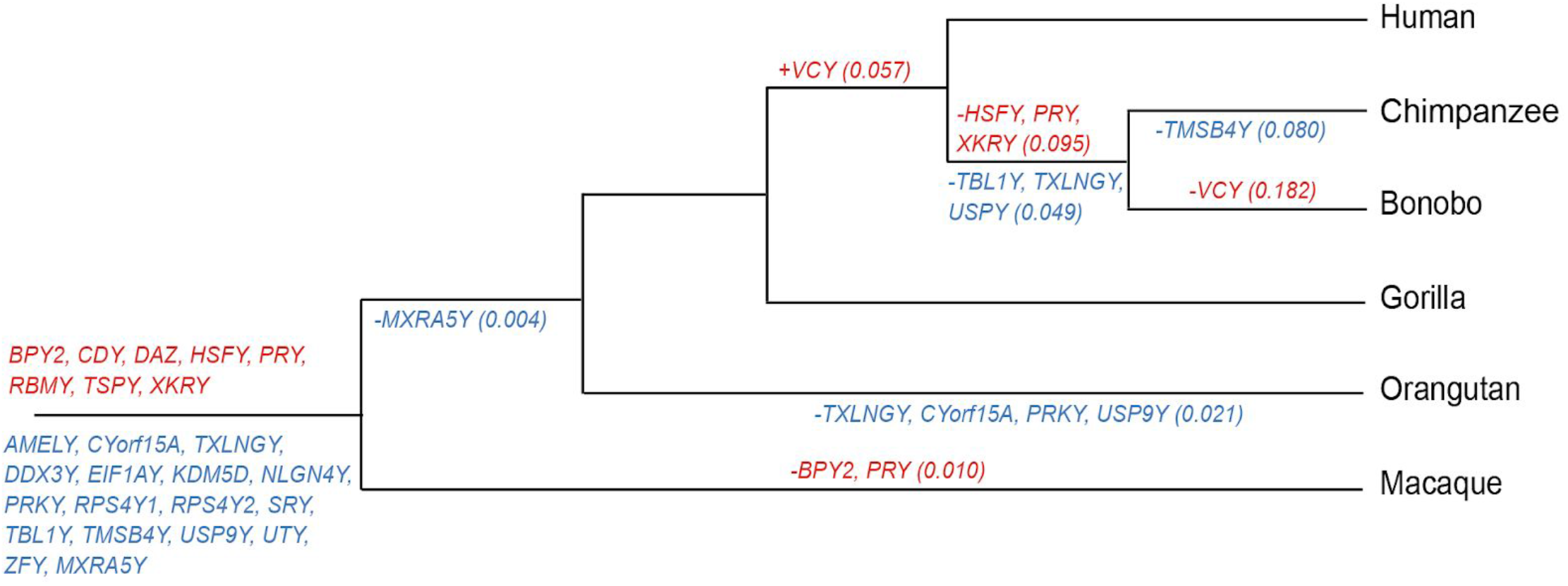
Evolution of Y chromosome gene content in great apes. The reconstructed history of gene birth and death for X-degenerate (blue) and ampliconic (red) genes was overlaid on the great ape phylogenetic tree (not drawn to scale), using macaque as an outgroup. The rates of gene birth and death (in events per million years) are shown in parentheses (for complete data see **Fig. S4**). The list at the root includes the genes that were present in the common ancestor of great apes and macaque. In addition to most of the genes on the human Y, the macaque Y harbors the X-degenerate *MXRA5Y* gene, which we found to be deleted in orangutan and pseudogenized in bonobo, chimpanzee, gorilla, and human. We currently cannot find a full-length copy of the *VCY* gene in bonobo. *TXLNGY* and *DDX3Y* are also known as *CYorf15B* and *DBY*, respectively.

### Conservation of human and chimpanzee palindrome sequences

Did the palindromes now present on the human Y (P1-P8) and chimpanzee Y (C1-C19) evolve before or after the great ape lineages split? To answer this question, we identified the proportions of human and chimpanzee palindrome sequences that aligned to bonobo, orangutan and gorilla Ys in our multi-species alignments (**Fig. 3A**, **Table S5**). Among human palindromes, P5 and P6 were the most conserved (covered by 89-97% of other great ape Y assemblies), whereas the majority of P3 sequences were human-specific (covered by only 31-38% of other great ape Y assemblies). Nevertheless, the common ancestor of great apes likely already had substantial lengths of sequences homologous to P1, P2, and P4-P8, and some sequences of P3 (**Fig. 3B**). Chimpanzee palindromes C17, C18, and C19 are homologous to human palindromes P8, P7, and P6, respectively (7). Therefore below we focused on the other chimpanzee palindromes and, following (7), divided them into five homologous groups: C1 (C1+C6+C8+C10+C14+C16), C2 (C2+C11+C15), C3 (C3+C12), C4 (C4+C13), and C5 (C5+C7+C9) (**Table S6**). The palindromes in the C3, C4, and C5 groups had substantial proportions (usually 70-94%) of their sequences covered by alignments with other great ape Ys (**Fig. 3A**). In contrast, most of C2 sequences (85%) were shared with bonobo, and a substantial proportion of C1 sequences was chimpanzee-specific. Nonetheless, the common ancestor of great apes likely already had large amounts of sequences homologous to group C3, C4, and C5 palindromes, and also some sequences homologous to group C1 and C2 palindromes (**Fig. 3B**).

**Figure 3.**
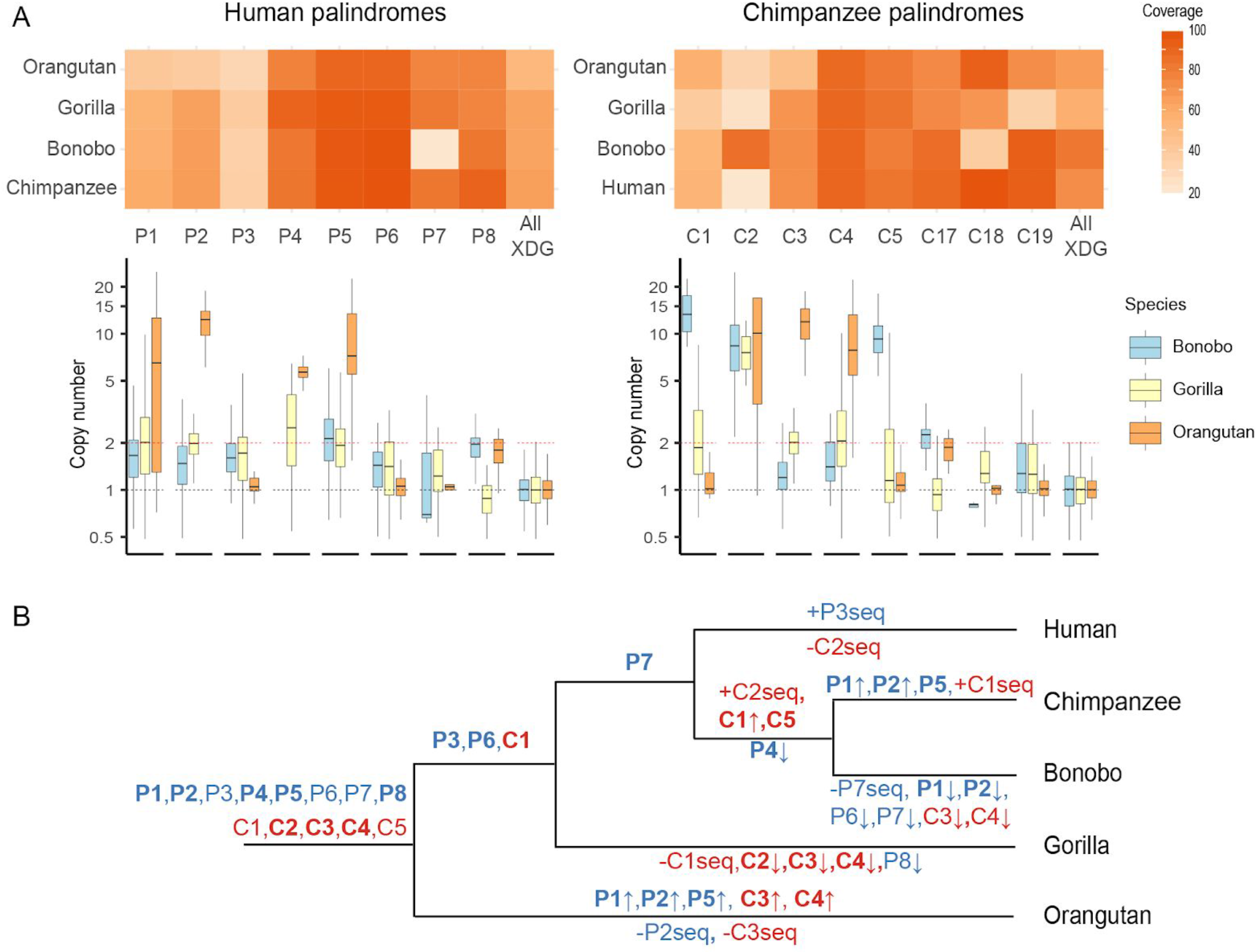
Evolution of sequences homologous to human and chimpanzee palindromes. **(A)** Heatmaps showing coverage for each palindrome in each species in the multi-species alignment, and box plots representing copy number (natural log) of 1-kb windows which have homology with human or chimpanzee palindromes. (**B**) The great ape phylogenetic tree (not drawn to scale) with evolution of human (shown in blue) and chimpanzee (purple) palindromic sequences overlayed on it. Palindrome names in bold indicate that their sequences were present in ≥2 copies. Negative (−) and positive (+) signs indicate gain and loss of palindrome sequence (possibly only partial), respectively. Arrows represent gain (↑) or loss (↓) of palindrome copy number. If several equally parsimonious scenarios were possible, we conservatively assumed a later date of acquisition of the multi-copy state for a palindrome (**Note S4**).

To determine whether the bonobo, orangutan, and gorilla sequences homologous to human or chimpanzee palindromes were multi-copy (i.e. present in more than one copy), and thus could form palindromes, in the common ancestor of great apes, we obtained their read depths from whole-genome sequencing of their respective males (**Fig. 3A**; see Methods). This approach was used because we expect that some palindromes were collapsed in our Y assemblies. We also used the data on the homology between human and chimpanzee palindromes summarized from the literature (6–8) (**Table S7**). Using maximum parsimony reconstruction, we concluded (**Note S4**) that sequences homologous to P4, P5, P8, C4, and partial sequences homologous to P1, P2, C2, and C3 were multi-copy in the common ancestor of great apes (**Fig. 3B**). Sequences homologous to P3, P6, and C1 were multi-copy in the human-gorilla common ancestor, those homologous to P7 were multi-copy in the human-chimpanzee common ancestor, and those homologous to C5 were multi-copy in the bonobo-chimpanzee common ancestor (**Fig. 3B**; **Note S4**).

### Species-specific multi-copy sequences in bonobo, gorilla, and orangutan

In addition to finding sequences homologous to human and/or chimpanzee palindromes, we detected 9.5 Mb, 1.6 Mb, and 3.5 Mb of species-specific sequences in our bonobo, gorilla, and orangutan Y assemblies (**Fig. S5**). By mapping male whole-genome sequencing reads to these sequences (see Methods), we found that 81%, 43%, and 32% of them in bonobo, gorilla, and orangutan had copy number of 2 or above (**Table S8**). Thus, large portions of Y species-specific sequences are multi-copy and might harbor species-specific palindromes.

### What drives conservation of gene-free palindromes P6 and P7?

Palindromes are hypothesized to evolve to allow ampliconic genes to withstand high mutation rates on the Y via gene conversion in the absence of interchromosomal recombination (10, 11). Two human palindromes—P6 and P7—do not harbor any genes, however the large proportions of their sequences are present and are multi-copy in most great ape species we examined (**Fig. 3A, Table S7**). We hypothesized that conservation of P6 and P7 might be explained by their role in regulation of gene expression. Using ENCODE data (22), we identified candidate open chromatin and protein-binding sites in P6 and P7 (**Fig. S7**). In P6, we found markers for open chromatin (DNase I hypersensitive sites) and histone modifications H3K4me1 and H3K27ac, associated with enhancers (23), in human umbilical vein endothelial cells (HUVEC). In P7, we found markers for CAMP-responsive element binding protein 1 (CREB-1), participating in transcription regulation (24), in a human liver cancer cell line HepG2. Interestingly, we did not identify any open-chromatin or enhancer marks in P6 and P7 in testis, suggesting that the sites we found above regulate genes expressed outside of this tissue.

### Frequent chromatin interactions between and within palindromes

Because Y ampliconic regions undergo Y-Y gene conversion and Non-Allelic Homologous Recombination (NAHR) (11), we hypothesized that these processes are facilitated by increased chromatin interactions. To evaluate this, we studied chromatin interactions on the Y utilizing a statistical approach specifically developed for handling Hi-C data originating from repetitive sequences (25). We used publicly available Hi-C data generated for human and chimpanzee induced pluripotent stem cells (iPSCs) (26) and for HUVEC (27). We found prominent chromatin contacts both between and within palindromes located inside ampliconic regions on the Y (**Fig. 4A-B**). In fact, the contacts in the human palindromic regions were significantly overrepresented when compared with the expectation based on the proportion of the Y occupied by palindromes (*p*<0.001, permutation test with palindromic/non-palindromic group categories; **Table S9**, **Fig. S8**), suggesting biological importance. Notably, we observed similar patterns for two different human cell types, as well as for both human and chimpanzee iPSCs (**Fig. 4A-B, Fig. S9**).

**Figure 4.**
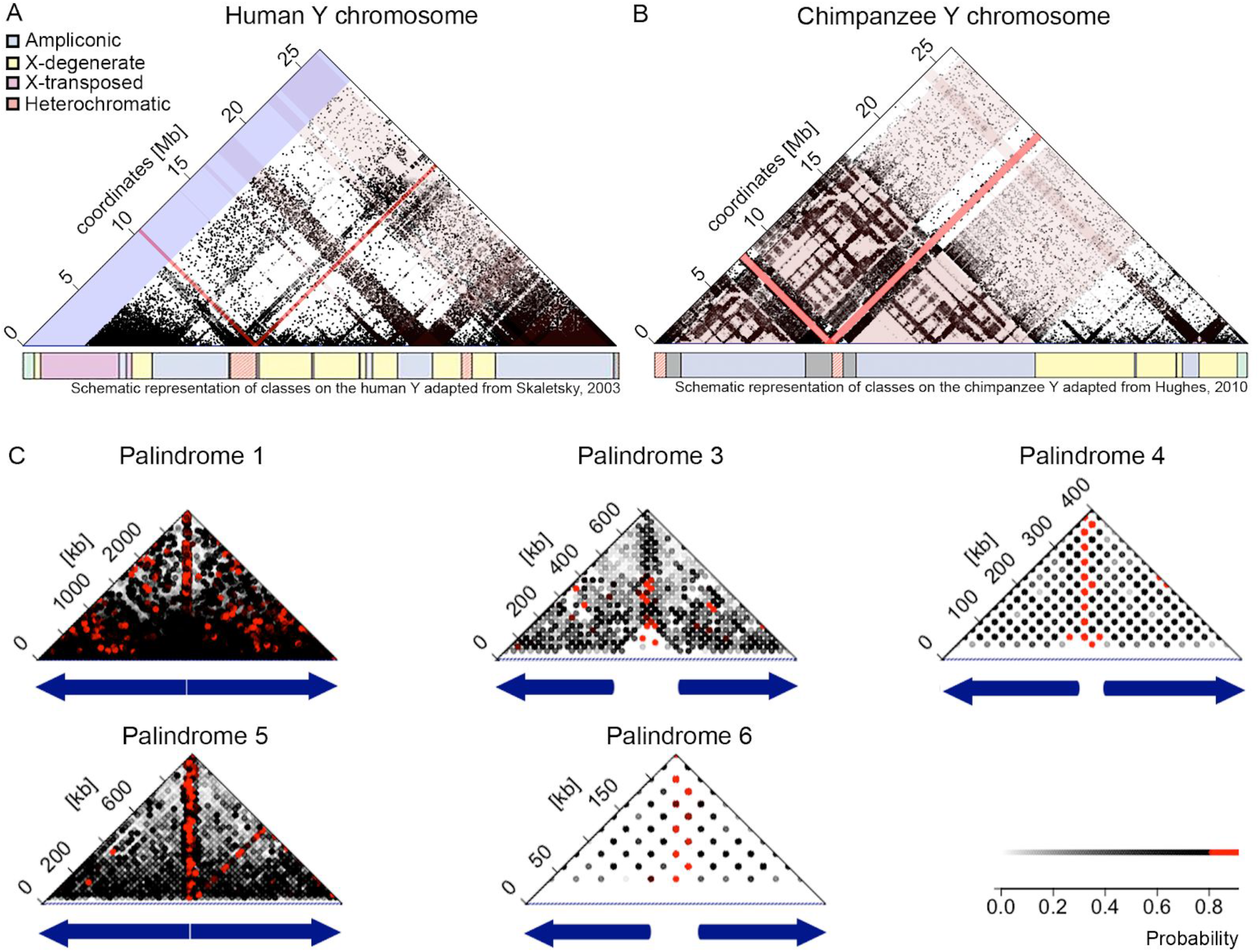
Chromatin contacts on the human and chimpanzee Y chromosomes, as evaluated from iPSCs. **A.** Human Y chromosome contacts with palindromes (highlighted in pink), pseudoautosomal regions (blue), and centromere (red). The schematic representation of the sequencing classes on the Y chromosome is adapted from (6). **B.** Chimpanzee Y chromosome contacts with palindromes (highlighted in pink). The schematic representation of the sequencing classes on the Y chromosome is adapted from (7). **C**. Chromatin interactions for the five largest palindromes on the human Y. To resolve ambiguity due to multi-mapping reads, each interaction was assigned a probability based on the fraction of reads supporting it (see Supplemental Methods for details). Palindrome arms are shown as blue arrows and the spacer is shown as white space between them.

We also hypothesized that arms of the same palindrome interact with each other via chromatin contacts. Our analysis of human Hi-C data from iPSCs (26) suggests that palindrome arms are indeed co-localized—a pattern particularly prominent for the large palindromes P1 and P5 (**Fig. 4C**). These results suggest that, in addition to the enrichment in the local interactions expected to be present in the Hi-C data (28), homologous regions of the two arms of a palindrome interact with each other with high frequency.

## Discussion

### Substitution rates

Higher substitution rates on the Y than on the autosomes, which we found across the great ape phylogeny, confirm another study (29) and are consistent with male mutation bias likely caused by a higher number of cell divisions in the male than the female germline (17). Higher autosomal substitution rates we detected in the *Pan* than *Homo* lineage corroborate yet another study (30) and can be explained by shorter generation time in *Pan*. A higher Y-to-autosomal substitution ratio (i.e. stronger male mutation bias) in the *Pan* than *Homo* lineage, as observed by us here, could be due to several reasons. First, species with sperm competition produce more sperm and thus undergo a greater number of replication rounds, generating more mutations on the Y and potentially leading to stronger male mutation bias, than species without sperm competition (17). Consistent with this expectation, chimpanzee and bonobo experience sperm competition and exhibit strong male mutation bias, as compared with no sperm competition (31) and weak male mutation bias in human and gorilla (**Note S2**). Contradicting this expectation, orangutans have limited sperm competition (31), but exhibit strong male mutation bias (**Note S2**). *Second*, a shorter spermatogenic cycle can increase the number of replication rounds per time unit and can elevate Y substitution rates, leading to stronger male mutation bias. In agreement with this explanation, the spermatogenic cycle is shorter in chimpanzee than in human (32, 33); the data are limited for other great apes. *Third*, a stronger male mutation bias would be expected in *Pan* than in *Homo* if the ratio of male-to-female generation times were respectively higher (34). However, the opposite is true: this ratio is higher in *Homo* than in *Pan* (34).

Phylogenetic studies produce estimates of male mutation bias that might be affected by ancient genetic polymorphism in closely related species (29). Even though we corrected for this effect (**Note S2**), our results should be taken with caution because of incomplete data on the sizes of ancestral great ape populations (35). Pedigree studies inferring male mutation bias are unaffected by ancient genetic polymorphism. One such study detected significantly higher male mutation bias in chimpanzee than in human (36), in agreement with our results, while another study found no significant differences in male mutation bias among great apes (37). These two studies analyzed only a handful of trios per species, and thus their conclusions should be re-evaluated in larger studies.

### Ampliconic sequences

We found that substantial portions of most human palindromes, and of most chimpanzee palindrome groups, were likely multi-copy (and thus potentially palindromic) in the common ancestor of great apes, suggesting conservation over >13 MY. Moreover, two of the three rhesus macaque palindromes are conserved with human palindromes P4 and P5 (20), indicating conservation over >25 MY. Our study also found species-specific amplification or loss of ampliconic sequences, indicating that their evolution is rapid. Thus, repetitive sequences constitute a biologically significant component of great ape Y chromosomes and their multi-copy state might be selected for.

Ampliconic sequences are thought to have evolved multiple times in diverse species to enable intrachromosomal, i.e. Y-Y, gene conversion (reviewed in (38)). Y-Y gene conversion can compensate for the degeneration in the absence of interchromosomal recombination on the Y by removing deleterious mutations (39, 40), can decrease the drift-driven loss of less mutated alleles, can lead to concerted evolution of repeats (11), and can increase the fixation rate of beneficial mutations (38). Yet, despite its critical importance for the Y, how Y-Y gene conversion occurs mechanistically is not well understood. Our analysis of Hi-C data suggested that ampliconic sequences and palindrome arms co-localize on the Y in both human and chimpanzee, potentially facilitating both Y-Y gene conversion and NAHR. The latter process is frequently used to explain rapid evolution of the ampliconic gene families’ copy number (41), as well as structural rearrangements (42), some of which lead to spermatogenic failure, sex reversal, and Turner syndrome (43).

Previous studies (e.g., reviewed in (10, 11, 38)) focused on the role of Y-Y gene conversion in preserving Y ampliconic gene families, which are critical for spermatogenesis and fertility (6), and suggested that this phenomenon explains the major adaptive role of palindromic sequences. Our findings suggest that conservation of some palindromes is driven not by spermatogenesis-related genes, but by regions regulating the expression of genes transcribed outside of testis and thus likely located outside of the Y. This observation should be examined in more detail in the future, but can potentially shift a paradigm in our understanding of Y chromosome functions. Indeed, our results imply that, in addition to carrying genes important for spermatogenesis, the Y chromosome participates in regulating gene expression in the genome. This echoes findings in *Drosophila* (e.g., (44)) and in the mouse Y chromosome, which contains a small, gene-free region that interacts with the rest of the genome (45).

### Gene content evolution

We inferred that the gene content in the common ancestor of great apes likely was the same as is currently found in gorilla, and included eight ampliconic and 16 X-degenerate genes (**Fig. 2**). Analyzing the data on ampliconic gene content (**Fig. 2**), palindrome sequence (**Fig. 3B**), and ampliconic gene copy number (Fig. 2 in (19)) evolution jointly, we can infer which ampliconic genes were present in the multi-copy state in the great ape common ancestor. Our results suggest that such ancestor had multi-copy sequences homologous to P1, P2, P4, P5 and P8 (**Fig. 3B**), which in human carry *DAZ, BPY2, CDY, HSFY, XKRY*, and *VCY* (5). Except for *VCY*, which was acquired by the human-chimpanzee common ancestor, the remaining five genes were likely present as multi-copy gene families in the common ancestor of great apes, because three of them (*DAZ*, *BPY2*, and *CDY)* are present as multi-copy in all great ape species (19), and the other two *(HSFY*and *XKRY)—*in all great ape species but chimpanzee and bonobo (19), in which they were lost (**Fig. 2**).

We discovered that there is only one gene family that was born across the whole great ape Y phylogeny— *VCY* was acquired by the common ancestor of human and chimpanzee. As a result, except for this branch, we found uniformly low rates of gene birth. A low rate of *ampliconic* gene birth contradicts predictions of high birth rate made in previous studies for such genes (38), but suggests that great ape radiation does not provide sufficient time for gene acquisition by ampliconic regions. Ampliconic regions on the Y chromosomes of several other mammals acquired such genes (46, 47), however the timing of such acquisitions is unknown.

We expected to observe a high death rate for X-degenerate genes, but a low death rate for ampliconic genes, because the former genes do not undergo Y-Y gene conversion and thus should accumulate deleterious mutations, whereas the latter genes are multi-copy and can be rescued by Y-Y gene conversion. Unexpectedly, the rates of gene death were similar between ampliconic and X-degenerate genes. Indeed, ~44.4% of ampliconic gene families were either deleted or pseudogenized, as compared with also ~43.8% of X-degenerate genes, across the great ape Y phylogenetic tree. While our data did not support our hypothesis, other findings suggest that death of ampliconic genes is a gradual process. Indeed, ampliconic gene families dead in some great ape species have reduced copy number in other species (19, 41), lowering the chances for Y-Y gene conversion. Thus, such genes are on the way to become non-essential and are at death’s door in great apes.

The rates of gene death varied among great ape species. In particular, we observed high rates of death in the lineages of bonobo, chimpanzee, and their common ancestor. What could be the evolutionary forces driving such a high rate of gene death, likely operating in the *Pan* lineage continuously since its divergence from the human lineage? First, gene-disrupting or gene deletion mutations could be hitchhiking in haplotypes with beneficial mutations. Positive selection might be acting in the *Pan* lineage due to sperm competition. No gene deaths in the human and gorilla lineages, experiencing no sperm competition, and low gene death rates in orangutan, experiencing limited sperm competition, are consistent with this explanation. Our tests for positive selection acting at protein-coding genes did not produce significant results, however indicated significantly elevated nonsynonymous-to-synonymous rate ratios for five genes in bonobo, chimpanzee, and/or their common ancestor (**Note S5**). We might have limited power to detect positive selection from phylogenetic data collected for closely related species. Second, the *Pan* Y could have undergone stronger drift leading to fixation of variants lacking genes that were already in the process of becoming non-essential. Predominant dispersal by females, and the associated low male effective population size, observed in bonobo and chimpanzee (48), as compared with predominant dispersal by males in gorilla and orangutan (3, 49), are consistent with this explanation.

### Future directions

Future studies should include sequencing of the Y chromosome for multiple individuals per species, and such data are expected to provide a resolution between the evolutionary scenarios driving high substitution and gene death rates in the *Pan* lineage. Future investigations should also focus on deciphering the sequences of different copies and isoforms of ampliconic genes (50), which should allow one to examine natural selection potentially operating at them in more detail. Because chromatin organization depends on the tissue of origin (27), the high prevalence of intra-ampliconic contacts we found in two somatic tissues should be confirmed in testis and sperm. Additionally, comparing chromatin organization and evolution of palindromes on the Y vs. X chromosomes should aid in understanding the unique role repetitive regions might play on the Y.

From a more applied perspective, the bonobo and orangutan Y assemblies presented here are useful for developing genetic markers to track male dispersal in these endangered species. This is of utmost importance because both species experience population decreases due to habitat loss. Therefore our results are expected to be of great utility to conservation genetics efforts aimed at restoring these populations.

## Materials and Methods

See Supplemental Methods for details.

### Assemblies and alignments

For bonobo and Sumatran orangutan, we generated and assembled (51) deep-coverage short sequencing reads from male individuals, and identified putative Y contigs by mapping them against the corresponding female reference assemblies (52). These contigs were then scaffolded with mate-pair reads (53). The orangutan Y assembly was further improved by merging (54) with another high-quality assembly generated with 10×Genomics technology (55). The bonobo Y assembly was improved by additional scaffolding with long Y-enriched Pacific Biosciences reads (56, 57). We improved the continuity of the gorilla Y assembly by merging two previously published assemblies (8, 58). To remove PARs, we filtered each species-specific Y assembly against the corresponding female reference genomes. Great ape Y assemblies were aligned with PROGRESSIVECACTUS (59). Substitution rates were estimated for alignments blocks containing all five species with the GTR model (60) implemented in PHYLOFIT (61).

### Gene content analysis

To retrieve the bonobo and orangutan genes, we aligned the scaffolds from their Y chromosome assemblies to the respective species-specific or closest-species-specific reference coding sequences using BWA-MEM (62). Novel gene predictions were evaluated with AUGUSTUS (63). The evolutionary history of Y gene content and gene birth and death rates was reconstructed using (21).

### Palindrome analysis

To analyze conservation of human and chimpanzee palindromes, we found all multi-species alignment blocks that overlap their coordinates and identified the percentage of non-repetitive bases in such blocks per species. To evaluate the copy number of sequences homologous to human and chimpanzee palindromes in bonobo, gorilla, and orangutan, we mapped whole-genome male sequencing reads to the corresponding 1-kb windows which overlap intervals of the human and chimpanzee Y palindromes using BWA-MEM (62) and compared their read depth with that of single-copy X-degenerate genes. The copy number of species-specific ampliconic sequences in bonobo and orangutan were evaluated similarly. Regulatory factor binding sites in human palindromes P6 and P7 were extracted from ENCODE (22). To analyze a potential enrichment of ampliconic interactions, Hi-C data (26, 27) were processed with MHI-C (25).

### Data and code availability

Sequencing data, assemblies and alignments are available under BioProject PRJNA602326. Code is available at https://github.com/makovalab-psu/great-ape-Y-evolution.

## Acknowledgments

We are grateful to L. Carrel, O. Ryder, M. Ferguson-Smith, S. Warris, K. Sahlin, F. Chiaromonte, A. Kenney, and the Smithsonian Institution for their assistance. This research was supported by NIH R01GM130691 and NSF DBI-1356529, IIS-1453527, and CCF-1439057.

