## Supplemental notes, tables, and figures for "Dynamic evolution of great ape Y chromosomes"

### Supplement to Cechova, Vegesna, et al.

**Supplement to Cechova, Vegesna, et al.** **1**

Supplemental Methods 3

Short- and long-read data to generate Y chromosome assemblies 3

Multi-species alignments 4

Substitution rate analysis 5

Gene content analysis 5

Palindrome analysis 6

Supplemental Notes 8

Supplemental Note S1. Developing a classifier and evaluating it. 8

Supplemental Note S2. Male mutation bias. 10

Supplemental Note S3. AUGUSTUS gene predictions 11

Supplemental Note S4. Evolutionary scenarios for palindromes. 13

Supplemental Note S5. Analysis of X-Y gene conversion and selection 15

Supplemental Tables 18

Table S1. The statistics for de novo bonobo, gorilla, and orangutan Y chromosome assemblies and for the human and chimpanzee reference Y chromosomes (1, 2). 18

Table S2. Alignment statistics. 19

Table S3. The estimated number of substitutions (after correcting for multiple hits). 20

Table S4. Gene birth and death rates (in events per millions of years) on the Y chromosome of great apes, as predicted using the Iwasaki and Takagi gene reconstruction model (33). 21

Table S5. The sequence coverage (percentage) of human palindromes P1-8 (arms) by non-human great ape Y assemblies. 22

Table S6. The sequence coverage (percentage) of chimpanzee palindromes C1-19 (arms) across great apes. 23

Table S7. The copy number for sequences homologous to (A) human and (B) chimpanzee palindromes. 24

Table S8. Summary of Y chromosome species-specific sequences. 25

Table S9. The number of observed chromatin contacts, followed by the chromatin interactions weighted by their probability (multi-mapping reads are allocated probabilistically (46)). The density of chromatin interactions is higher in palindromes. The table is based on human iPSC data (47). See also Fig. S8. 26

Table S10. The generated-here and previously unpublished sequencing datasets used for assembly generation and classification, including their summary information. 27

Table S11. The number of variants before and after the polishing step. 28

Table S12. The coordinates of palindromes on panTro6 and hg38 Y chromosome. 29

Table S13. The list of ENCODE datasets analyzed in the search for regulatory elements in P6 and P7. 30

Supplemental Figures 31

Figure S1. Flowcharts for the assemblies of (A) bonobo, (B) Sumatran orangutan, and (C) gorilla. 31

Figure S2A. Sequence comparisons between human (hg38) and bonobo, gorilla and orangutan Y chromosome assemblies. 34

Figure S2B. Sequence comparisons between chimpanzee (panTro6) and bonobo, gorilla and Sumatran orangutan Y chromosome assemblies. 35

Figure S3. Protein-coding gene sequence retrieval in the new Y assemblies. 36

Figure S4. (A, C, E) Thresholds used for classification of windows into X-degenerate versus ampliconic, and (B, D, F) average copy number for overlapping 5-kb windows. 38

Figure S5. Shared and lineage-specific sequences in multi-species alignments. 39

Figure S6. Reconstructed gene content of great apes. 40

Figure S7. IGV screen shots of peaks of DNase-seq, H3K4me1 and H3K27ac marks on human palindrome P6, and of CREB1 on human palindrome P7. 41

Figure S8. Chromatin interactions on the human Y chromosome. 42

Figure S9. Hi-C contact map generated for Human Umbilical Vein Cells (HUVEC), using the data from (49). 43

References 44

### Supplemental Methods

#### Short- and long-read data to generate Y chromosome assemblies

We used human and chimpanzee Y sequences from version hg38 and panTro6 assemblies, respectively (1, 2). In both cases, PARs were masked (we used human coordinates 10,001-2,781,479 and 56,887,903-57,217,415 for PAR1 and PAR2, respectively, as provided by ENSEMBL (https://useast.ensembl.org/info/genome/genebuild/human_PARS.html) and 26,208,347-26,341,346 for PAR in chimpanzee, identified in house using LASTZ (3) alignments). For the gorilla Y chromosome, there are two published assemblies: the one produced from short- and long-read data by our group (4), and the one recently generated using PacBio reads pre-processed with PACASUS to remove within-read duplications (5). We merged the two assemblies to obtain a more continuous and complete version; the merging was performed with METASSEMBLER v.1.5 (6). Since METASSEMBLER additionally uses information from mate-pair reads, we used the previously-generated GY19 library (4) and set insert sizes to range from 298 to 6,444 bp in the configuration file.

For both orangutan and bonobo (**Fig. S1**), we first generated short, paired-end reads for a male individual. Sumatran orangutan male (ID 1991-51) genomic DNA provided by the Smithsonian Institution was extracted from testis using DNeasy Blood and Tissue Kit (Qiagen). Bonobo male (ID PR00251) genomic DNA was extracted using the same kit from the fibroblast cell line provided by the San Diego Zoological Society. Illumina paired-end PCR-free libraries with insert size of ~1,000 bp were constructed from both DNA samples and sequenced on HiSeq2500 using 251-bp paired-end sequencing protocol (approximately 600 million reads were generated for each library). The reads were assembled using DISCOVAR (7). To identify putative Y-chromosomal contigs, we processed each of these two male assemblies with DISCOVERY (8) using the male-to-female abundance threshold of 0.4 and using "-abundance-min 4". The resulting contigs were scaffolded with filtered mate-pair 250-bp reads, generated for the same individuals using Nextera Mate Pair Sample Preparation Kit (with median insert size of 8 kb), using BESST v.2.2.8 (9). Filtering of mate pairs was performed as follows. All mate-pair reads were mapped to the respective (bonobo or orangutan) whole-genome male DISCOVAR assembly using BWA-MEM v.0.7.17-r1188 (10). From these alignments, reads putatively originating from Y chromosome were retained and used for scaffolding. The next step, aimed at further improving the continuity of Y assemblies, differed between orangutan and bonobo. The orangutan Y assembly was merged (using METASSEMBLER v.1.5 (6)) with an assembly generated for orangutan male AG06213 with 10×Genomics (**Table S10**). To aid METASSEMBLER, one can provide mate-pair reads; we used mate-pair reads from individual 1991-51 and set the insert size range from 5,088 to 12,683 bp in the configuration file. High-molecular-weight genomic DNA from Sumatran orangutan male (ID AG06213) was extracted from fibroblast cell line (provided by L. Carrel, PSU) using MagAttract HMW DNA kit (Qiagen) and used to construct a 10x Genomics library. The resulting reads were assembled with SUPERNOVA assembler v.1.0.2 (11). The bonobo Y assembly was augmented with PACASUS-corrected (default parameters) (5) flow-sorted Y-enriched PacBio reads from a bonobo male Ppa_MFS using SSPACE-LONGREAD (v.1-1) (12), and additional scaffolding and assembly were performed with PBJELLY (PBSuite_14.9.9, parameters <blasr>-minMatch 8 -minPctIdentity 85 -bestn 1 -nCandidates 20 -maxScore -500 -nproc 6 -noSplitSubreads</blasr>) (13).

The subsequent processing steps were the same for the orangutan Y and bonobo Y assemblies. For the orangutan, we first called variants from the first male (ID 1991-0051) relative to the assembly we obtained after merging with 10×Genomics data. The variants were called with FREEBAYES (v.1.3.1) (14) using settings "--ploidy 1 --min-alternate-fraction 0.6", and filtered for quality (biopet-vcffilter --minQualScore 30). We then used BCFTOOLS to 'polish' the assembly by consensus calling (BCFTOOLS consensus -H 1, **Table S11**). Using the same approach, we polished the bonobo Y assembly using the variants from bonobo male ID PR00251.

To remove the pseudoautosomal region (PAR) and potential residual autosomal contamination, all scaffolds in gorilla, bonobo and orangutan Y assemblies were filtered against corresponding gorilla (gorGor4), bonobo (panPan2), and orangutan (ponAbe3) female reference genomes. We mapped the assembly scaffolds to the female reference using BLASR v.5.3.3 (15) with parameters: "blasr --bestn 1 --nproc 32 --hitPolicy randombest --header --minPctSimilarity 95". Scaffolds that had at least 95% sequence identity to the female genome were excluded from the assemblies.

#### Multi-species alignments

To align the Y chromosomes of five great ape species, several alignment algorithms were evaluated: SIBELIAZ (16), MULTIZ-TBA (17), MULTIZ-ROAST (17), and PROGRESSIVECACTUS (18). The alignments were evaluated both by comparing the proportion of aligned base pairs and visually, i.e. by comparing the assembled coverage in non-human great ape species sequences aligned to the human X-degenerate genes. PROGRESSIVECACTUS showed the most promise and was chosen for all further analyses. We soft-masked repeats prior to the alignment of assemblies using REPEATMASKER (19) with the following settings: *RepeatMasker -pa 63 -xsmall -species Primates ${assembly}.rmsk.fa* (19). To generate multiple sequence alignment with PROGRESSIVECACTUS, we used default parameters and did not provide a guide tree. The alignment was converted from hal (20) to MAF format using HAL2MAF (parameterized with, for example, --noAncestors --refGenome hg_Y --maxRefGap 100 --maxBlockLen 10000). The role of the MAF reference sequence was given to each species in turn, to create a separate MAF file for each, as well as for the PROGRESSIVECACTUS-inferred ancestral sequence.

We split the MAF file by species subset, producing a separate MAF file containing all blocks that only involve the particular species in that subset. The MAF file of blocks containing all five species was converted to MULTI-FASTA by MAF to FASTA tool, run at https://usegalaxy.org/ (GALAXY version 1.0.1), using an option "One Sequence per Species". Sequence identity was calculated from the MULTI-FASTA file using the custom script multi_fasta_to_pairwise_identity.py.

Independently, sequence identity distributions were derived from pairwise alignments as follows. For each pair of species (s1, s2) LASTZ (3) was used to align s1's Y assembly and s2's. Masking was disabled, allowing alignment of duplicated elements. Substitution scores were identical to those used by us in Tomaszkiewicz et al. (4). The exact LASTZ command line was: "lastz s1.chrY[unmask] s2.chrY[unmask] W=12 Q=human_primate.scores." Alignments were then post-processed as follows. For each base in s1, the highest identity alignment covering that base was chosen, and base counts were collected for identities at 0.1% intervals. Average sequence identity was computed from these binned counts, excluding any unaligned bases.

Percentage of sequence aligned in multiple alignment was computed as follows. For the portion of species s1 aligning to s2, all MAF blocks containing s1 and s2 (as well as, possibly, other species) were extracted. These were reduced to the aligning intervals in s1 by computing the mathematical union of all s1 intervals in any of the extracted blocks. The combined length of these intervals is the number of bases in s1 that have an alignment to s2. This length was divided by the number of non-N bases in s1.

Percentage of sequence aligned in pairwise alignments was computed as follows. For the portion of species s1 aligning to s2, the s1-to-s2 LASTZ alignment was reduced to the aligning intervals in s1 by computing the mathematical union of all s1-intervals in any alignment block. The combined length of these intervals is the number of aligning bases in s1. This length was divided by the number of non-N bases in s1.

#### Substitution rate analysis

To estimate the substitution rates on the Y chromosome, we used the ancestor-based five-species alignment described in the previous section. To provide conservative estimates of substitution rates, we removed duplicates from the alignment using MAFDUPLICATEFILTER from the MAFTOOLS suite (version 0.1) (21), similar to (22). This step replaces multiple sequences from the same species by the sequence closest to the consensus of an alignment block. We also converted all alignments blocks to the positive strand of the ancestral sequence (maf_flip_for_ref.py). Again, to obtain conservative estimates, we only retained alignment blocks where all five species were present, thus largely restricting our analysis to X-degenerate regions. This step was performed using script *parse_cactus* (available on GITHUB) and the python libraries *bx.align.maf* and *bx.align.tools*. The final filtered alignment was then used to pick the best-fitted substitution model using JMODELTEST (23, 24). Taking into account AIC and BIC of the various models tested with JMODELTEST and the availability of models implemented in PHYLOFIT (25), we decided to proceed with GTR (also called REV) model (26) with variable substitution rates (--nrates=4). Using this model and our filtered alignment, we ran PHYLOFIT (25) with the following settings: phyloFit -E -Z -D 42 --subst-mod REV --nrates 4 --tree "(((hg_Y,panTro_Y),gorGor_Y),ponAbe_Y)". The estimated branch lengths were used for all the downstream analysis. The output was then visualized using R script and library APE (27)(28).

#### Gene content analysis

**Analysis of genes homologous to human Y genes.** To retrieve the bonobo and orangutan Y chromosome X-degenerate and ampliconic genes, we aligned the scaffolds from bonobo and Sumatran orangutan Y chromosome assemblies to species-specific or closest-species-specific reference coding sequences using BWA-MEM (v.0.7.10) (10). Next, we visualized the alignment results in Integrative Genomics Viewer (IGV) (v.2.3.72), and protein-coding consensus sequences were retrieved and checked for ORFs. However information about *RPS4Y2* and *MXRA5Y* remained missing. We used gene predictions (see next paragraph) and testis-specific transcriptome assemblies (29) from the same publication to study the presence of the *RPS4Y2* and *MXRA5Y* genes (**Fig. S6**).

**Predicting novel genes in bonobo and orangutan assemblies.** We used AUGUSTUS (30) (--species=human --softmasking=on --codingseq=on) to predict genes in the Y chromosome assemblies of bonobo and orangutan. From the list of predicted genes we retained those that have a start and stop codon. Using BLASTP from BLAST (2.9.0; -db uniprot_sprot.pep -max_target_seqs 1 -evalue 1e-5 -num_threads 10) (31) we annotated the predicted gene sequences. Based on BLAST annotation, we classified all the genes which are not annotated by BLASTP as candidate *de novo* genes. From these genes, we retained those that had >90% sequence identity to the sequence in the UniProt (32) protein database and were covered by at least 90% of the gene sequence in the BLAST output. To make sure that the predicted genes are on the Y chromosome and do not represent misassemblies, we performed an extra filtering step where we used only those genes which are present on contigs which align to either human or chimpanzee Y chromosomes. The resulting gene annotations which are not found on the human Y chromosome were assigned as candidate novel genes translocated to the Y. The longest predicted transcript sequences for genes annotated as Y-chromosome-specific were obtained without the above filters for validation of missing gene content in bonobo and orangutan. The gene predictions which were annotated as X homologs (*TXLNG*, *PRKX*, *NLGNX*, etc.) of Y chromosome genes were also classified as Y genes.

**Reconstruction of gene content evolution in great apes.** Once we obtained the complete Y chromosome gene content in great apes, we converted the gene content into binary values which represent the presence or absence of a gene in a species. The complete deletion or pseudogenization of gene/gene families is represented by ‘zero’ and the presence of an uninterrupted gene is represented by ‘one’. Using the model developed by Iwasaki and Takagi (33), we reconstructed the evolutionary history of (separately ampliconic and X-degenerate) genes across great apes. The phylogenetic tree of great apes (34), along with the table representing the presence or absence of genes (**Fig. S6**), was used as an input to generate the rate of gene birth and gene death for each branch of the tree, and the reconstructed gene content at each internal node of the tree. To obtain the rate in the units of events per million years, the rate of gene birth, as well as the rate of gene death, was divided by the length of the branches (in millions years) (33).

#### Palindrome analysis

**Human and chimpanzee palindrome arm sequence conservation.** The PROGRESSIVECACTUS output file was converted to MAF (see the preceding paragraph) twice. The first time we set --refGenome to 'human', which results in a human-centric MAF file (where the first line represents sequences from the human Y) and the second time we set --refGenome parameter to 'chimpanzee' to obtain a chimpanzee-based MAF file. The coordinates of the human and chimpanzee palindromes (PanTro4) were obtained from a previous publication (4). Chimpanzee palindrome coordinates were converted to the PanTro6 version using the LIFTOVER utility from the UCSC Genome Browser (35). For few chimpanzee palindromes for which LIFTOVER failed to convert the coordinates, we generated a dotplot of chimpanzee Y to itself using GEPARD (36) and identified the locations of palindromes from the dotplot. To overcome redundancy of sequence between the arms of palindrome, we further obtained the coordinates of the first arm of the palindrome for human (**Table S12A**) and chimpanzee (**Table S12B**), from which the palindrome coverage was calculated.

With the use of ALIGNIO from BIOPYTHON (37), we parsed the MAF files to obtain all the alignment blocks which overlapped given palindrome arm coordinates for each species and summed the number of sites covered by these parsed alignment blocks. For each alignment block within the palindrome arm, we considered only those sequences which had less than 5% gaps and counted the number of aligned nucleotides per species at sites where the palindrome arm is unmasked (softmasked with REPEATMASKER (19)). Next we calculated the number of nonrepetitive characters in the whole palindrome arm and used it to calculate the percentage of non-repetitive alignment within the palindrome arm per species.

**Palindrome sequence read depth in bonobo, gorilla, and orangutan.** We used a pre-established pipeline used to identify human and chimpanzee palindrome sequences in Y assemblies (4). The bonobo, gorilla, and orangutan Y contigs were broken into non-overlapping 1-kb windows and each window was aligned to human (hg38) and (separately) chimpanzee (panTro6) Y chromosome sequences using LASTZ (version 1.04.00) (--scores=human_primate.q, --seed=match12, --markend) (3) separately. The substitution scores were set identical to those used for published LASTZ alignments of primates (38). The best alignment for each window was retrieved from the LASTZ output. All the windows with alignments of >800 bp were considered as homologs to human or chimpanzee Y palindromes and used in the downstream analysis.

The whole-genome male sequencing reads from orangutan, gorilla, and bonobo (**Table S10**) were mapped to the human and (separately) chimpanzee Ys using BWA-MEM (version 0.7.17-r1188)(10) and the resulting output files were sorted and indexed using samtools (version 1.9) (39). Using COMPUTEGCBIAS and CORRECTGCBIAS functions from DEEPTOOLS (version 3.3.0) (40), we performed GC correction of the sequencing read depths. Finally, using BEDTOOLS (v2.27.1)(41) 'coverage' function, we obtained the read depth of the windows homologous to human and chimpanzee palindrome sequences (of whole palindromes) identified as described in the previous paragraph. We compared the read depths of palindromic sequence windows to that of windows overlapping X-degenerate genes.

**Finding the copy number for ampliconic regions that are species-specific**. From the multiple sequence alignment of the Y chromosomes of great apes (species-centric), we extracted the alignment blocks which were unique to a particular species and not aligned to the other Y chromosomes. We filtered the species-specific blocks which were >100 bp in size and obtained their GC-corrected read depth from male whole-genome sequencing data. To calculate the copy number of the resulting blocks we divided their read depth by the median read depth of windows which overlap X-degenerate genes from the palindrome copy number analysis (read depth of single-copy regions).

**Search for regulatory factor binding sites in human palindromes P6 and P7.** We checked for the presence of sites specific to DNA binding proteins on human palindromes P6 and P7, which could imply the presence of functionality related to gene regulation. Initially we used the ENCODE track (http://genome.ucsc.edu/ENCODE/) at the UCSC Genome Browser (35) to search for epigenetic modifications in the palindrome regions (42). In particular, we used the track from the Bernstein Lab at the Broad Institute containing H3K27Ac and H3K4Me1 data on seven cell lines from ENCODE, from which we identified human umbilical vein endothelial cells (HUVEC) to have signals in palindrome P6. Later we extended our search to ENCODE data portal (https://www.encodeproject.org/) from where we downloaded the BAM files to look for the presence of peaks (**Table S13**)(43). We used the "Search by region" page under Data in the ENCODE data portal (https://www.encodeproject.org/region-search/) and GRCh38 setting to search for files which are related to the coordinates of palindromes P6 (chrY:16159590-16425757) and P7 (chrY:15874906-15904894). The resulting files were visualized in the UCSC Genome Browser to identify signals in the peaks, and from this information we identified signals in human liver cancer cell line HepG2 for P7. The current ENCODE data processing pipeline by default filters out reads with low mapping quality (i.e. mapping in multiple places of the genome), as a result we did not find any peaks/signals in the majority of the datasets (https://www.encodeproject.org/pipelines/). To overcome this limitation, we manually downloaded the unfiltered BAM files (HUVEC: H3K27Ac, H3K4Me1 and DNase-seq; Testis:H3K27Ac, H3K4Me1 and Dnase-seq; and HepG2: CREB1; Table S13), which include low-mapping-quality reads. We performed peak calling using MACS2 (version2.1.4; -f BAM --broad --broad-cutoff 0.05) (44) and the control samples were used whenever available. We used integrative genomics viewer (IGV) (version 2.4.19)(45) to visualize the data.

**Hi-C analysis.** We used MHI-C (46) to process the Hi-C data generated by (47). Both human and chimpanzee samples originated from pluripotent stem cells (iPSCs), processed with protocol using *Mbo I* restriction enzyme (with the GATCGATC recognition site) and sequenced on Illumina Hi-Seq 4000. The sequencing reads were downloaded from SRA (accessions SRR1658709 and SRR8187262 for human and chimpanzee, respectively) and subsampled to the first 100 million reads. The original MHI-C scripts were modified to match the reference genome and other settings; the resolution was set to 10,000, cutsite was set to 0, and normalization method was set to KR (48) with uniPrior (built based on unique bin pairs within valid interaction distance, see https://github.com/keleslab/mHiC). Additionally, we analyzed a biological sample extracted from umbilical vein endothelial cells of a newborn male, cultured in cell line HUVEC (CC-2517), further subjected to the *in situ* Hi-C protocol with *Mbo I* restriction enzyme and sequenced on Illumina HiSeq 2000 (49). For this analysis, we used the first 50 million reads of SRR1658570 dataset.

In order to calculate the enrichment of ampliconic interactions, we defined these regions separately in human and chimpanzee. In human, we used the corresponding coordinates of eight palindromes (4). Because chimpanzee Y is composed of a higher number of intertwined amplicons and 19 palindromes, the continous ampliconic part spanning the *q* arm of this chromosome, as well as a large part of *p* arm adjacent to the centromere, was analyzed as a single unit.

### Supplemental Notes

#### Supplemental Note S1. Developing a classifier and evaluating it.

We developed a classifier to determine which assembly scaffolds are ampliconic and which are X-degenerate. Ideally, Y ampliconic regions would have a higher copy number in the reference genome than X-degenerate regions, however, Y ampliconic regions are often collapsed in next-generation-sequencing-based assemblies (50). Our classifier therefore combines the copy count in the reference with mapping read depth information, since collapsed Y ampliconic regions will have a higher number of mapping reads than single-copy X-degenerate regions, in whole-genome-sequencing datasets originating from male individuals. We proceeded to classify scaffolds in our orangutan, bonobo, and gorilla Y assemblies (such annotations are already available in the human and chimpanzee Y assemblies (1, 2)). To test classification performance, we examined scaffolds carrying known X-degenerate and ampliconic genes (Table SN1A). The classification was successful in the vast majority of cases (accuracy was 85%, 100%, and 88% for bonobo, gorilla, and orangutan, respectively, Table SN1A).

**Table SN1A.** The number of X-degenerate and ampliconic genes that were confidently mapped to scaffolds classified as X-degenerate or ampliconic, per our classifier.

| Species | Gene type | Classified as X-degenerate | Classified as ampliconic |
|---|---|---|---|
| GORILLA | ampliconic | 2 | **5** |
| | X-degenerate | **12** | 1 |
| BONOBO | ampliconic | 1* | **5** |
| | X-degenerate | **9** | 0 |
| ORANGUTAN | ampliconic | 1 | **2** |
| | X-degenerate | **5** | 0 |

*BPY is present in a single copy in gorilla

##### Methods

To classify scaffolds as either ampliconic or X-degenerate (PAR regions have been filtered out in our assemblies), each assembly was divided into 5-kb windows with 2-kb overlaps. First, for each window, we calculated copy number using a modified version of AMPLICONE (51) that can handle scaffolds instead of a single continuous reference. AMPLICONE takes into account mappability (calculated using the GEM mappability program (52) for k=101), repetitive element content as provided by REPEATMASKER (Open-4.0.7, RepBaseRepeatMaskerEdition-20181026), and performs GC correction. X-degenerate regions are expected to have similar copy number estimates (i.e. estimates based on sequencing depth), whereas copy number estimates for ampliconic regions are expected to be higher and vary greatly, depending on the copy number of the underlying amplicons in the genome and in the assembly. Fully resolved amplicons, such as palindrome arms, are present multiple times in the assembly, and thus each window carrying them maps equally well to multiple places in the assembly. Thus, all windows were mapped back to the assembly and, after excluding secondary alignments, all multi-mapping windows were flagged as potentially ampliconic (windows with MAPQ=0 as produced by BWA-MEM (10)). If a window was both nonrepetitive (non-zero copy number by AMPLICONE) and multi-mapping, it was assigned as ampliconic. Additionally, all uniquely mapping windows (MAPQ=60) with high copy number for species-specific thresholds (see Table SN1B) were also assigned as ampliconic. Uniquely mapping windows (MAPQ=60) with copy number below the species-specific thresholds were assigned as X-degenerate. This led to the majority of windows classified as either ampliconic or X-degenerate. Two types of windows remained unassigned; those that map uniquely but contain no copy-number information from AMPLICONE, and those that contain copy-number information but have mapping quality below 60. All such windows had their assignment interpolated. As these windows were internally represented as missing values (NA), we used na.approx function from the R package zoo (53). Each scaffold was then classified as either ampliconic or X-degenerate based on the majority vote of all underlying windows.  A small subset of our assemblies (0.12, 0.33, and 0.71 Mb in bonobo, gorilla and orangutan) remained unclassified.

**Table NS1B.** The copy-number thresholds for X-degenerate (<) and ampliconic (≥) windows, the size of the classified ampliconic and X-degenerate regions, and the corresponding number of scaffolds.

| | Gene type | threshold | size [Mb] | # scaffolds |
|---|---|---|---|---|
| GORILLA | ampliconic | | 3.95 | 71 |
| | X-degenerate | 0.95 | 10.02 | 161 |
| BONOBO | ampliconic | | 10.8 | 2,521 |
| | X-degenerate | 0.5 | 12.5 | 967 |
| ORANGUTAN | ampliconic | | 2.2 | 670 |
| | X-degenerate | 0.75 | 14.5 | 358 |

For validation, we mapped coding sequences of X-degenerate and ampliconic genes from chimpanzee, gorilla, and Sumatran orangutan to verify that the scaffolds carrying these genes were classified correctly. We required strict mapping to avoid false hits; first we used blat (v. 36) with default parameters and then required at least thirty matches (matches≥30), the recovery of at least 20% of the original coding sequence query ((matches/qSize)≥0.2), and 99% of matching bases ((matches/(matches+misMatches))≥0.99). All accompanying scripts are available from https://bitbucket.org/biomonika/assembly_classification/ and in-house scripts *run_copy_number.sh* and evaluate.sh that output the annotation of scaffolds as a .gff file.

### Supplemental Note S2. Male mutation bias.

Using branch-specific values for autosomal and Y-chromosomal substitution rates from Fig. 1, we obtained the following values of α, or the male-to-female substitution rate ratio (54):

| Species | Substitution rate on the Y | Substitution rate on autosomes (A) | Y/A | α |
|---|---|---|---|---|
| **Human** | 0.0098 | 0.0066 | 1.48 | 2.88 |
| **Chimp** | 0.0037 | 0.0021 | 1.76 | 7.40 |
| **Bonobo** | 0.0044 | 0.0025 | 1.76 | 7.33 |
| **Gorilla** | 0.0155 | 0.0089 | 1.74 | 6.74 |
| **Orang** | 0.0582 | 0.0268 | 2.17 | ∞ |
| **BC** | 0.0088 | 0.0046 | 1.91 | 22.0 |
| **BCH** | 0.0046 | 0.002 | 2.30 | ∞ |

The species-specific estimates of α we obtained above follow the trend observed in our previous study (see Table 1 in (55)) based on a comparison of substitution rates at a much shorter genetic region (~10 kb) homologous between chromosomes Y and 3. Namely, we also observe that α is lower in human than in bonobo or gorilla. Note that our estimates are derived from closely related species and thus might be inaccurate because of ancient genetic polymorphism (55). This phenomenon is difficult to correct for in branch-specific estimates. However, it can be accounted for in pairwise estimates.

Focusing on pairwise comparisons we presented in the Results, we computed α using both uncorrected and corrected by ancestral polymorphism autosomal rate estimates. In primates, the Y chromosome has much lower diversity than autosomes do (56), and thus we did not correct for it. The results are shown below:

| | Y | A | Y/A | α | Corrected A* | Corrected Y/A | Corrected α |
|---|---|---|---|---|---|---|---|
| **Gorilla-human** | 0.0299 | 0.0175 | 1.71 | 5.86 | 0.01592 | 1.88 | 15.41 |
| **Gorilla-chimp** | 0.0326 | 0.0176 | 1.85 | 12.54 | 0.01602 | 2.03 | ∞ |
| **Gorilla-bonobo** | 0.0333 | 0.0180 | 1.85 | 12.33 | 0.01642 | 2.03 | ∞ |

*We subtracted 0.00158 -- the diversity estimated from gorilla populations (57) -- from our autosomal substitution rate estimates.

Again, consistent with the data presented in Results, we observed larger differences between the Y chromosomal and autosomal substitution rate estimates for gorilla-chimpanzee and gorilla-bonobo comparisons, than for the gorilla-human comparisons.

We also evaluated the potential effect of ancient genetic polymorphism on the ratio of pairwise estimates of autosomal substitution rates we present in Results. We found that this effect is minimal.

| | Y | A uncorrected | A ratio | A corrected* | Corrected ratio to gorilla-human comparison |
|---|---|---|---|---|---|
| **Gorilla-human** | 0.0299 | 0.0175 | | 0.0159 | |
| **Gorilla-chimp** | 0.0326 | 0.0176 | 1.006 | 0.0160 | 1.0063 |
| **Gorilla-bonobo** | 0.0333 | 0.018 | 1.029 | 0.0164 | 1.0314 |

*We subtracted 0.00158 - the diversity estimated from gorilla populations (57) -- from the autosomal substitution rate.

#### Supplemental Note S3. AUGUSTUS gene predictions

AUGUSTUS (30) predicted 219 genes on the bonobo Y assembly, of which 25 complete or partial genes represent homologs of known human protein-coding genes. In the case of Sumatran orangutan, AUGUSTUS predicted 90 genes of which 33 complete or partial genes represent homologs of known human protein-coding genes. After implementing requirements of gene predictions (1) to have start and stop codons, and (2) be present on contigs that align to human or chimpanzee Y, we did not find any novel genes on the Y chromosome of orangutan, however we found two candidates—*SUZ12* and *PSMA6*—which have >95% identity and >90% coverage to the gene homologs on the autosomes of bonobo.

A possible transposition of the autosomal *SUZ12* gene (located on human chromosome 17) onto the bonobo Y chromosome was predicted (Table SN3) based on the limited number of introns (one intron), in contrast to its autosomal homolog, which has 15 introns (NM_015355). The *SUZ12* gene has no matches to human and gorilla Y, however the first 121 bp of its predicted sequence align to chimpanzee (palindromes C2, C11 and C15) and orangutan Y. However, when we aligned testis RNA-seq data to the predicted *SUZ12* gene on the bonobo Y chromosome, the first exon with the start codon was not expressed (Fig. SN3A), whereas the second exon was expressed (Fig. SN3B). The single nucleotide variants in the RNA-seq reads mapping to the second exon are consistent with the variants of the *SUZ12* gene present on chromosome 17. Thus, we concluded that the translocated *SUZ12* was pseudogenized on bonobo Y.

The *PSMA6* gene was also predicted in bonobo, which shared 99.6% identity with its homolog on the chimpanzee Y and 97.8% identity on human Y. However, there were no homologous sequences in the orangutan and gorilla Y assemblies. The *PSMA6* gene on human Y was annotated as a pseudogene in the Entrez database (Gene ID: 5687). Therefore, we concluded that *PSMA6* is also a pseudogene on the bonobo. Thus, no novel genes, as compared to the human Y chromosome genes, were found on the bonobo and orangutan Y chromosomes.

**Table SN3. Gene annotation of the *SUZ12* homolog on the bonobo Y, as predicted by AUGUSTUS**

| Sequence name | Feature | Start | End | Strand |
|---|---|---|---|---|
| Contig591 | gene | 74 | 61572 | - |
| Contig591 | transcript | 74 | 61572 | - |
| Contig591 | stop_codon | 74 | 76 | - |
| Contig591 | CDS | 74 | 2353 | - |
| Contig591 | CDS | 61453 | 61572 | - |
| Contig591 | start_codon | 61570 | 61572 | - |

**Figure SN3A. The IGV** (45) **view of the first exon of the SUZ12 homolog on the bonobo Y.** Testis-specific RNA-seq reads (SRA ID: SRR306837) were mapped to bonobo Y assembly using BWA-MEM (10).

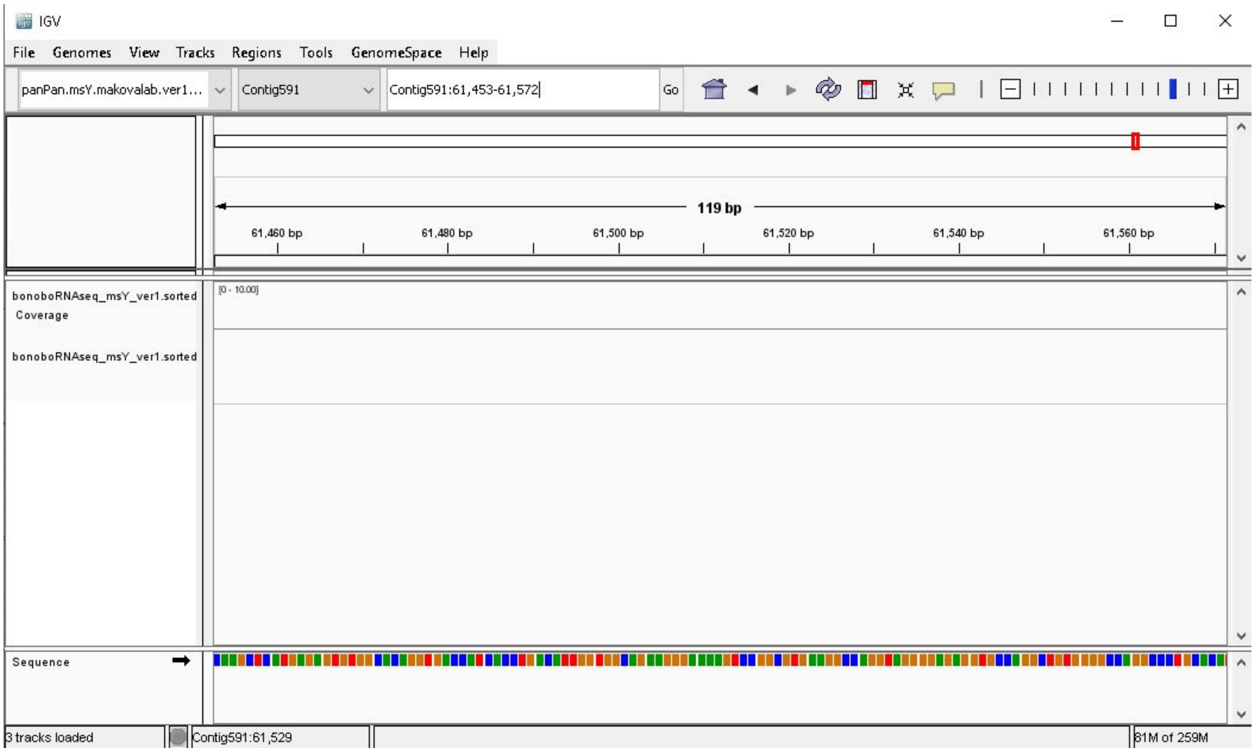

**Figure SN3B. IGV view of second exon on SUZ12 homolog on bonobo Y.** Testis specific RNASeq reads (SRA ID: SRR306837) were mapped to bonobo Y assembly using BWA-MEM (10).

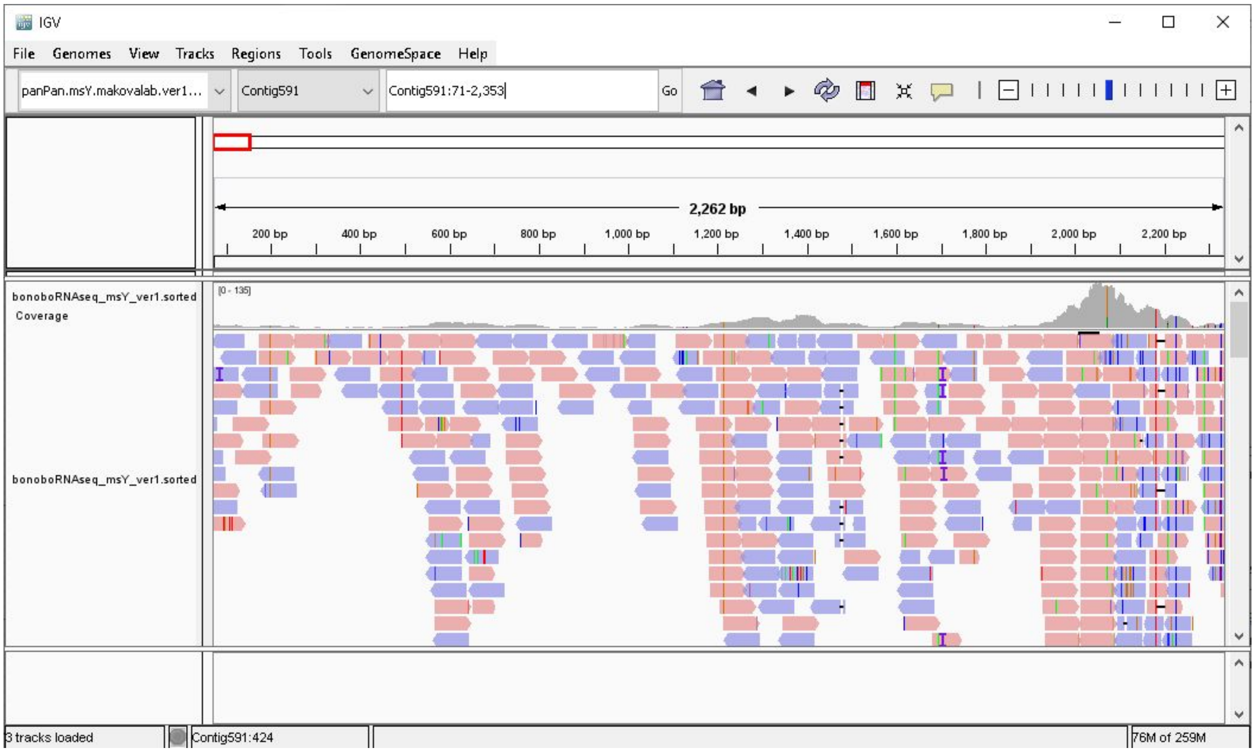

#### Supplemental Note S4. Evolutionary scenarios for palindromes.

**Palindrome 4.** We used two different approaches to obtain information about the conservation of human and chimpanzee palindromes across great apes. First, we used the multiple sequence alignment of Y chromosome assemblies to obtain the coverage of these palindromes. In the case of P4 we observed 23,175 bp of alignment in the bonobo assembly (**Table S5**). This analysis gave us the percentage of P4 present in bonobo Y assembly, however it did not infer that there is a continuous 23-kb block of palindrome P4 on bonobo Y. P4 could be highly fragmented due to Y chromosome degradation or rearrangements and the multiple sequence alignment can still capture such homologous fragments of the palindromes. The longest blocks in the Y multiple sequence alignment that overlap with P4 are 2-4 kb in length and these alignment blocks included sequences from gorilla and human Ys. In the remaining species, sequences homologous to P4 are mostly represented by gaps. Second, to identify the copy number of P4 homologs present in other species, we used alignments generated with LASTZ (3) based alignments. Non-overlapping 1-kb windows of the assembly were aligned to human P4 using LASTZ. The read depth of windows with >80% identity to P4 was used to estimate the copy number of P4. However, we did not find any 1-kb window in bonobo Y which maps to human palindrome P4 with >80% identify. Since we did not find high-confidence windows aligning to P4 in bonobo, we concluded that it is highly fragmented in this species.

**Evolution of sequences homologous to human and chimpanzee palindromes.** *Common ancestor of great apes.* Partial sequences of all human and chimpanzee palindromes were present. P1, P2, P4, P5, P8, C2, C3, C4, and C17 were in multi-copy state (Tables SN4A-B). P3, P6, and C1 might have been in single- or multi-copy state (Tables SN4A-B).

*Orangutan.* Increase in copy number of P1,P2, P5, C3, and C4 (Fig. 3B, Table SN4A-B). Loss of C3 (≈25% loss in coverage Table S6) and P2 (≈27% loss in coverage Table S5) segments when compared to other great apes.

*Common ancestor of gorilla, human, bonobo, and chimpanzee.* P3, P6, and C1 are in a multi-copy state either at this node or in the common ancestor of great apes (Tables SN4A-B).

*Gorilla.* Loss of copy number for P8, C2, and C17 compared to bonobo and orangutan (Fig. 3, Tables SN4A-B). C3 and C4 had more than two copies in human, chimpanzee and orangutan, however only two copies in gorilla (Tables SN4A-B). For C2, bonobo and orangutan have higher copy number when compared to gorilla. Loss of segments in C1 (≈15% loss in coverage) and C19 (≈40-60% loss in coverage) when compared to other great apes (Table S6).

*Common ancestor of human, bonobo, and chimpanzee.* All the palindromes are assumed to be multi-copy with the exception of C5, which could have been in a single-copy state (Tables SN4A-B). Gorilla and orangutan have more than two copies of P4 which is in two copies or lost in human, bonobo and chimpanzees (Tables SN4A-B).

*Human.* Palindrome P3 has ≈30-35% covered in other great apes (Table S5), so we assume that the remaining portion of P3 is human-specific. Humans also lost most of the sequences homologous to palindrome C2; we observe some sequence homologous to C2 on the human Y, however they are degraded and not visible on an alignment in human and chimpanzee Y dot plot (1). Therefore, we assume C2 was deleted human.

*Common ancestor of bonobo and chimpanzee.* Gain of C2 segment, both bonobo and chimpanzee share 85% coverage whereas the other great apes cover 20-30% of C2, which implies a *Pan* genus specific gain of C2 sequences (Table S6). C1 and C5 groups are present in more than two copies in both chimpanzee (1) and bonobo, where as other species have two or fewer copies of these palindromes (Fig. 3A). The *Pan* genus lost P4, we observe that sequence homologous to P4 are present in bonobo and chimpanzee Y, however they are degraded and not visible as an alignment in the human and chimpanzee Y dot plot (1). Therefore, we assume that P4 was deleted.

*Bonobo*. Bonobo lost copies of P1, P2, P6, P7, C3, C4, C18, and C19 (Fig. 3A, Table SN4A-B). It also experienced loss of segments in C18 (≈30-60% loss in coverage Table S6) and P7 (≈60% loss in coverage Table S5) when compared to other great apes.

*Chimpanzee*. Chimpanzee gained copies of P1, P2, and P5 as these palindromes share homology with C3 and C4, which are in multi-copy in chimpanzee (1). Chimpanzee also gained a segment of C1, a palindrome which has <50% coverage with the other great ape Y chromosomes (Table S6).

**Table SN4A. Reconstructing human palindrome evolution using maximum parsimony.** The values from extant species were taken from Table S7 and rounded to the following numbers of copies: <1.34 => "1", "1.33-1.66" =>"1-2", "1.66-2.5" => "2", ">2.5" => "M" ("more than two").

| | P1 | P2 | P3# | P4 | P5 | P6 | P7 | P8 |
|---|---|---|---|---|---|---|---|---|
| **Bonobo** | 1-2 | 1-2 | 1-2 | 0 | 2 | 1-2 | 1 | 2 |
| **Chimpanzee** | M | M | 1$ | 0 | M | 2 | 2 | 2 |
| **BC*** | 1-2-M | 1-2-M | 1 | 0 | 2-M | 2 | 1-2 | 2 |
| **Human** | 2 | 2 | 2 | 2 | 2 | 2 | 2 | 2 |
| **BCH**** | 2 | 2 | 1-2 | 2 | 2 | 2 | 2 | 2 |
| **Gorilla** | 2 | 2 | 2 | 2 | 2 | 1-2 | 1 | 1 |
| **BCHG***** | 2 | 2 | 2 | 2 | 2 | 2 | 1-2 | 1-2 |
| **Orangutan** | M | M | 1 | M | M | 1 | 1 | 2 |
| **GA****** | 2-M | 2-M | 1-2 | 2-M | 2-M | 1-2 | 1 | 2 |

*Bonobo-chimpanzee common ancestor
**Bonobo-chimpanzee-human common ancestor
***Bonobo-chimpanzee-human-gorilla common ancestor
****Common ancestor of great apes
$We conservatively assigned the copy number of P3 as 1 in chimpanzee to make sure we do not inflate its combined copy number.
#We can conservatively assume that P3 became multi-copy in BCHG, but instead it might have been multi-copy in the common ancestor of great apes and lost its multi-copy status in orangutan instead

**Table SN4B. Reconstructing chimpanzee palindrome evolution using maximum parsimony.** The values from extant species were taken from Table S7 and rounded to the following numbers of copies: <1.34 => "1", "1.33-1.66" =>"1-2", "1.66-2.5" => "2", ">2.5" => "M" ("more than two").

| | C1 group | C2 group | C3 group | C4 group | C5 group | C17 | C18 | C19 |
|---|---|---|---|---|---|---|---|---|
| **Chimpanzee** | M | M | M | M | M | 2 | 2 | 2 |
| **Bonobo** | M | M | 1 | 1-2 | M | 2 | 1 | 1 |
| **BC*** | M | M | 1-M | M | M | 2 | 2 | 1-2 |
| **Human** | 2 | 0 | M | M | 1 | 2 | 2 | 2 |
| **BCH**** | 2-M | M | M | M | 1-M | 2 | 2 | 2 |
| **Gorilla** | 2 | M | 2 | 2 | 1 | 1 | 1 | 1 |
| **BCHG***** | 2 | M | 2-M | 2-M | 1 | 1-2 | 1 | 1-2 |
| **Orangutan** | 1 | M | M | M | 1 | 2 | 1 | 1 |
| **GA****** | 1-2 | M | M | M | 1 | 2 | 1 | 1 |

*Bonobo-chimpanzee common ancestor
**Bonobo-chimpanzee-human common ancestor
***Bonobo-chimpanzee-human-gorilla common ancestor
****Common ancestor of great apes

#### Supplemental Note S5. Analysis of X-Y gene conversion and selection

**Gene conversion.** Prior to analyzing selection, we had to perform an analysis of X-Y gene conversion, as this process can interfere with selection detection. We studied the incidence of X-Y gene conversion between 12 X-degenerate genes and their homologs on the X chromosome (from 16 X-degenerate genes present on the human Y we excluded *CYorf15A*, *CYorf15B*, *RPS4Y1*, and *RPS4Y2,* as parts of these genes have repeats on the Y and homologs on the X, making detection of gene conversion difficult). Gene conversion was examined using a multiple-sequence alignment of the X and Y chromosomes.

We softmasked the X and Y chromosomes of great apes using REPEATMASKER (19) (RepeatMasker -pa 63 -xsmall -species Primates ${assembly}.rmsk.fa). PROGRESSIVECACTUS (18) was used to align the chromosomes. A guide tree which pairs the X and Y chromosomes from the same species list was used (((chimpY, chimpX),(bonoboY, bonoboX),(humanY, humanX),(gorillaY, gorillaX),(sorangY, sorangX))). The resulting alignment output from PROGRESSIVECACTUS was converted to MAF format by HAL2MAF (20) using human Y as a reference (--noAncestors --refGenome humanY --maxRefGap 100 --maxBlockLen 10000). We then parsed the alignment blocks (retaining blocks longer than >50bp; range of gene conversion tracts we observed in human was 55-290 bp, as was observed in previous studies (58, 59)) which fall within the coordinates of X-degenerate genes. We did not perform additional filtering based on repeat content within the alignment blocks. For each block, the alignments which constitute the sequences from both the X and Y chromosomes for a species were collected in a FASTA file. We used GENCONV (60) ( /w9  /lp -nolog) to identify gene conversion events based on multiple sequence alignment files. By default, the output of GENCONV constitutes a global list of high-confidence gene conversion events after multiple testing for each sequence pair. We also used /lp parameters with which a second list of significant gene conversions for all possible pairwise comparisons GENCONV performed was generated. We used a p-value cutoff of 0.05, as GENCONV provides p-values after correcting for multiple comparisons. We used pairwise comparisons to address gene conversion in cases where a chromosome was represented by more than one sequence in the alignment. From the GENCONV output, we parsed events that constitute gene conversion between the X and Y chromosome from the same species and retained events which are longer than 50 bp.

In total, we detected 143 candidate gene conversion events up to 50-410 bp in length (minimal length of 50 bp was used for detection), including 46 high-confidence ones (Table S5NA). Among these, most events were observed in the genes from younger strata—30 in *PRKY* (stratum 5), nine in *NLGN4Y* (stratum 4), four in *AMELY* (stratum 4), and two in *TBL1Y* (stratum 4). Higher homology between these genes and their X homologs, as opposed to between genes from older strata and their X homologs, is expected to facilitate gene conversion, which work most efficiently with homology >92% (58). No gene conversion events >50 bp in size were found in the exonic regions of X-degenerate genes, thus this process should not affect our selection analysis.

**Table S5NA. Gene conversion between X and Y chromosomes of great apes using GENECONV.** The values represent the number of high-confidence gene conversion events with significant *p*-values (<0.05) which are corrected for multiple comparisons across all sequence pairs. The values in brackets represent the total number of gene conversion events with significant *p*-values (<0.05) for comparisons corrected for the length of the alignment.

| Gene (Stratum) | Bonobo | Chimpanzee | Human | Gorilla | Orangutan | Total |
|---|---|---|---|---|---|---|
| ***AMELY* (4)** | 1 (1) | 1 (1) | 1 (1) | 1 (1) | 0 | 4 (4) |
| ***DBY* (3)** | 0 (1) | 0 (1) | 0 | 0 | 0 (1) | 0 (3) |
| ***EIF1AY* (3)** | 0 | 0 | 0 | 0 | 0 | 0 |
| ***KDM5D* (2)** | 0 | 0 | 0 | 0 | 0 | 0 |

| | | | | | | |
|---|---|---|---|---|---|---|
| ***NLGN4Y* (4)** | 2 (6) | 2 (7) | 3 (7) | 1 (8) | 1 (3) | 9 (31) |
| ***PRKY* (5)** | 8 (25) | 10 (27) | 4 (17) | 8 (17) | 0 | 30 (86) |
| ***SRY* (1)** | 0 | 0 | 0 | 0 | 0 | 0 |
| ***TBL1Y* (4)** | 1 (4) | 0 (2) | 1 (7) | 0 (3) | 0 (2) | 2 (18) |
| ***TMSB4Y* (3)** | 0 | 0 | 0 | 0 | 0 | 0 |
| ***USP9Y* (3)** | 0 | 0 | 0 | 0 | 0 | 0 |
| ***UTY* (3)** | 0 | 0 | 0 | 0 | 0 | 0 |
| ***ZFY* (3)** | 0 | 0 | 0 | 0 | 1 (1) | 1(1) |
| **Total** | 12 (37) | 13 (38) | 9 (32) | 10 (29) | 2 (7) | 46(143) |

**Selection.** We used the CODEML module of PAML (version 4.8) (61) to detect branch-specific differences in the nonsynonymous-to-synonymous rate ratios and to test for positive selection acting on X-degenerate genes (excluding pseudogenes *CYorf15A* and *CYorf15B* (*TXLNGY*)) and ampliconic genes (excluding *VCY* present only in two species) in human, chimpanzee, bonobo, gorilla, Sumatran orangutan, Bornean orangutan, and macaque. Coding sequences of Y-chromosomal genes were retrieved from GENBANK or deciphered in this study and aligned using CLUSTALW (62). The phylogenies were generated with the Neighbor-Joining method (63) (with 1,000 bootstrap replicas) as implemented in MEGA7 (64). First, for each gene, the one-ratio model (assuming the same nonsynonymous-to-synonymous rate ratio $\omega$ for the entire tree) was compared with the two-ratio model (assuming that the branch-specific ratio $\omega_s$ is different from the background ratio $\omega_o$). When the difference between the two models was significant, this indicated that the synonymous-to-nonsynonymous rate ratio was different for the branch tested. In these cases, to test for positive selection, the model assuming the foreground ratio $\omega$ to be fixed at 1 (neutral evolution) was compared against an alternative model with branch-specific $\omega > 1$ (positive selection).

We found a total of eight gene-branch combinations that had foreground nonsynonymous-to-synonymous rate ratio significantly different than the background nonsynonymous-to-synonymous rate ratio (Table S5NB). We observed significantly different nonsynonymous-to-synonymous rate ratios on the chimpanzee and bonobo ancestor than other lineages for three X-degenerate genes (*DDX3Y*, *EIF1AY*, and *PRKY*) and one ampliconic gene (*CDY*). We also detected significantly different nonsynonymous-to-synonymous rate ratios in bonobo for two X-degenerate genes (*DDX3Y* and *EIF1AY*), in chimpanzee for *ZFY*, and in human for *RPS4Y2*. However, none of these ratios was significantly higher than one, providing no evidence for positive selection.

**Table S5NB. Gene-branch combinations with significantly higher branch-specific than background nonsynonymous-to-synonymous rate ratio.**

| Gene | Branch | Background $\omega_o$ | Branch-specific $\omega_s$ | P-value for testing $\omega_s > \omega_o$ | P-value for testing $\omega_s > 1$ |
|---|---|---|---|---|---|
| ***CDY*** | BC | 0.44 | 3.08 | 0.02 | 0.30 |
| ***DDX3Y*** | Bonobo | 0.19 | 1.11 | 0.02 | 0.54 |
| ***DDX3Y*** | BC* | 0.18 | 1.44 | 0.004 | 0.48 |

| | | | | | |
|---|---|---|---|---|---|
| ***EIF1AY*** | Bonobo | 0.02 | >1 (division by 0) | 0.03 | 0.45 |
| ***EIF1AY*** | BC | 0.02 | >1 (division by 0) | 0.03 | 0.44 |
| ***PRKY*** | BC | 0.18 | 1.64 | 0.03 | 0.11 |
| ***RPS4Y2*** | Human | 0.16 | >1 (division by 0) | 0.0002 | 1.00 |
| ***ZFY*** | Chimpanzee | 0.04 | >1 (division by 0) | 0.04 | 0.44 |

*BC: bonobo-chimpanzee common ancestor

### Supplemental Tables

Table S1. The statistics for *de novo* bonobo, gorilla, and orangutan Y chromosome assemblies and for the human and chimpanzee reference Y chromosomes (1, 2).

| Species | Assembly length (Mb) | NG50[1] (in bp, using G=8.5 Mb) | N50[2] (in bp) | Number of scaffolds | Ns (in Mb) |
|---|---|---|---|---|---|
| **Orangutan** | 17.4 | 1,388,499 | 773,523 | 1,178 | 1.3 |
| **Gorilla** | 14.3 | 150,017 | 95,534 | 268 | 0.008 |
| **Bonobo** | 23.4 | 153,556 | 32,114 | 3,590 | 0.8 |
| **Chimpanzee[1]** | 26.4 | - | - | 1 | 1.1 |
| **Human[2]** | 57.2 | - | - | 1 | 33.6 |

[1]NG50: the size of the scaffold for which half of the conserved X-degenerate regions (set to 8.5 Mb based on estimates in human (2)) is in scaffolds that are equal to or larger than this size.
[2]N50: the size of the scaffold for which half of the assembly is in scaffolds that are equal to or larger in size.

### Table S2. Alignment statistics.

**(A)** Portion of a species' Y chromosome assembly aligning to each other species, in PROGRESSIVECACTUS (18) multi-species alignments. Percentage shown is the portion of bases in the column-species Z that has any pairwise alignment to the row-species W. Denominator is the non-N count of the column-species Z. **(B)** Portion aligning to each other species, in LASTZ (3) pairwise alignments. **(C)** Average identity in PROGRESSIVECACTUS (18) alignment blocks containing all five species. **(D)** Average identity in LASTZ (3) pairwise alignments.

**A.** Portion of a species aligning to each other species, in **PROGRESSIVECACTUS multi-species** alignments.

| proportion of / aligning to | Human Y | Chimpanzee Y | Bonobo Y | Gorilla Y | Orangutan Y |
|---|---|---|---|---|---|
| Human Y | — | 77.22% | 47.52% | 75.72% | 62.89% |
| Chimpanzee Y | 66.39% | — | 52.69% | 66.86% | 60.65% |
| Bonobo Y | 61.74% | 82.98% | — | 64.77% | 57.76% |
| Gorilla Y | 57.79% | 58.47% | 36.76% | — | 52.51% |
| Orangutan Y | 54.93% | 63.26% | 38.51% | 61.20% | — |

**B.** Portion aligning to each other species, in **LASTZ pairwise** alignments.

| proportion of / aligning to | Human Y | Chimpanzee Y | Bonobo Y | Gorilla Y | Orangutan Y |
|---|---|---|---|---|---|
| Human Y | — | 92.13% | 64.59% | 89.61% | 86.40% |
| Chimpanzee Y | 84.26% | — | 65.58% | 85.19% | 85.85% |
| Bonobo Y | 84.05% | 95.99% | — | 86.50% | 85.34% |
| Gorilla Y | 78.21% | 83.21% | 56.05% | — | 79.45% |
| Orangutan Y | 75.53% | 83.57% | 55.67% | 79.44% | — |

**C.** Average identity in **PROGRESSIVECACTUS multi-species** alignment blocks containing all five species.

| identity of / aligning to | Human Y | Chimpanzee Y | Bonobo Y | Gorilla Y | Orangutan Y |
|---|---|---|---|---|---|
| Human Y | — | 97.90% | 97.85% | 97.20% | 93.72% |
| Chimpanzee Y | 97.89% | — | 99.22% | 96.98% | 93.58% |
| Bonobo Y | 97.80% | 99.17% | — | 96.90% | 93.47% |
| Gorilla Y | 97.20% | 96.99% | 96.95% | — | 93.55% |
| Orangutan Y | 93.59% | 93.45% | 93.37% | 93.42% | — |

**S2D.** Average identity in **LASTZ pairwise** alignments.

| identity of / aligning to | Human Y | Chimpanzee Y | Bonobo Y | Gorilla Y | Orangutan Y |
|---|---|---|---|---|---|
| Human Y | — | 95.76% | 95.58% | 95.81% | 92.47% |
| Chimpanzee Y | 95.60% | — | 97.84% | 94.53% | 91.95% |
| Bonobo Y | 95.19% | 98.27% | — | 94.30% | 91.83% |
| Gorilla Y | 95.01% | 93.99% | 93.98% | — | 91.85% |
| Orangutan Y | 92.46% | 91.91% | 92.28% | 92.71% | — |

#### Table S3. The estimated number of substitutions (after correcting for multiple hits).

(**A**) between gorilla and chimpanzee, and between gorilla and human; (**B**) between gorilla and bonobo, and between gorilla and human, considering autosomes and Y chromosome separately, with corresponding $\chi^2$ statistics and *p*-value showing an elevation in the *Pan* lineage. We used a test similar to the relative rate test used in (22). The last column shows the alignment length.

**A**

| | | | | | |
|---|---|---|---|---|---|
| **Autosomes** | **N subst. gorilla -> chimp** | **N subst. gorilla -> human** | **ratio** | **Difference from 1 $\chi^2$, *p*-value** | **Alignment length** |
| | 40,594,903 | 40,364,251 | 1.006 | 657.1; $<1\times10^{-5}$ | 2,306,528,605 bp |
| **Y** | **N subst. gorilla -> chimp** | **N subst. gorilla -> human** | **ratio** | **χ2** | **Alignment length** |
| | 154,937 | 142,105 | 1.090 | 554.5; $<1\times10^{-5}$ | 4,752,665 bp |

**B**

| | | | | | |
|---|---|---|---|---|---|
| **Autosomes** | **N subst. gorilla -> bonobo** | **N subst. gorilla -> human** | **ratio** | **χ2** | **Alignment length** |
| | 41,517,515 | 40,364,251 | 1.029 | 16,243.7; $<1\times10^{-5}$ | 2,306,528,605 bp |
| **Y** | **N subst. gorilla -> bonobo** | **N subst. gorilla -> human** | **ratio** | **χ2** | **Alignment length** |
| | 158,264 | 142,105 | 1.114 | 869.7; $<1\times10^{-5}$ | 4,752,665 bp |

Table S4. Gene birth and death rates (in events per millions of years) on the Y chromosome of great apes, as predicted using the Iwasaki and Takagi gene reconstruction model (33).

BC - common ancestor of bonobo and chimpanzee; BCH - common ancestor of bonobo, chimpanzee, and human; BCHG - common ancestor of bonobo, chimpanzee, human, and gorilla. GA - common ancestor of great apes.

| | X-degenerate genes | | Ampliconic genes | |
|---|---|---|---|---|
| **Branch** | **Birth rate** | **Death rate** | **Birth rate** | **Death rate** |
| **Bonobo-BC** | $1.00 \times 10^{-4}$ | $1.00 \times 10^{-4}$ | $1.00 \times 10^{-4}$ | 0.182 |
| **Chimpanzee-BC** | $1.00 \times 10^{-4}$ | $8.00 \times 10^{-2}$ | $1.00 \times 10^{-4}$ | $1.00 \times 10^{-4}$ |
| **Human-BCH** | $1.90 \times 10^{-5}$ | $1.90 \times 10^{-5}$ | $1.90 \times 10^{-5}$ | $1.90 \times 10^{-5}$ |
| **Gorilla-BCHG** | $1.43 \times 10^{-5}$ | $1.43 \times 10^{-5}$ | $1.43 \times 10^{-5}$ | $1.43 \times 10^{-5}$ |
| **Orangutan-GA** | $7.14 \times 10^{-6}$ | $2.06 \times 10^{-2}$ | $7.14 \times 10^{-6}$ | $7.14 \times 10^{-6}$ |
| **Macaque-Root** | $3.45 \times 10^{-6}$ | $3.45 \times 10^{-6}$ | $3.45 \times 10^{-6}$ | $9.92 \times 10^{-3}$ |
| **BC-BCH** | $2.35 \times 10^{-5}$ | $4.89 \times 10^{-2}$ | $2.35 \times 10^{-5}$ | $9.54 \times 10^{-2}$ |
| **BCH-BCHG** | $5.71 \times 10^{-5}$ | $5.71 \times 10^{-5}$ | $5.71 \times 10^{-2}$ | $5.71 \times 10^{-5}$ |
| **BCHG-GA** | $1.43 \times 10^{-5}$ | $1.43 \times 10^{-5}$ | $1.43 \times 10^{-5}$ | $1.43 \times 10^{-5}$ |
| **GA-Root** | $6.67 \times 10^{-6}$ | $4.04 \times 10^{-3}$ | $6.67 \times 10^{-6}$ | $6.67 \times 10^{-6}$ |

Table S5. The sequence coverage (percentage) of human palindromes P1-8 (arms) by non-human great ape Y assemblies.
The repeats annotated by REPEATMASKER (19) were excluded from the calculations.

| Human palindrome | Length, bp | | | | | Coverage, percentage | | | |
|---|---|---|---|---|---|---|---|---|---|
| | Chimpanzee | Bonobo | Gorilla | Orangutan | Human* | Chimpanzee | Bonobo | Gorilla | Orangutan |
| **P1** | 362895 | 340679 | 334030 | 249906 | 608650 | 59.62 | 55.97 | 54.88 | 41.06 |
| **P2** | 50379 | 50144 | 49092 | 29106 | 76387 | 65.95 | 65.64 | 64.27 | 38.10 |
| **P3** | 64460 | 62335 | 68254 | 55671 | 179793 | 35.85 | 34.67 | 37.96 | 30.96 |
| **P4** | 77345 | 76543 | 83948 | 72509 | 93979 | 82.30 | 81.45 | 89.33 | 77.15 |
| **P5** | 158548 | 158405 | 157040 | 148243 | 166929 | 94.98 | 94.89 | 94.08 | 88.81 |
| **P6** | 35555 | 35436 | 34535 | 32729 | 36832 | 96.53 | 96.21 | 93.76 | 88.86 |
| **P7** | 4439 | 1134 | 4385 | 4134 | 5414 | 81.99 | 20.95 | 80.99 | 76.36 |
| **P8** | 15027 | 13698 | 12688 | 12783 | 16728 | 89.83 | 81.89 | 75.85 | 76.42 |
| **Total** | 768648 | 738374 | 743972 | 605081 | 1184712 | 64.88 | 62.33 | 62.80 | 51.07 |

*Palindrome arm length

Table S6. The sequence coverage (percentage) of chimpanzee palindromes C1-19 (arms) across great apes.

The repeats annotated by REPEATMASKER (19) were excluded from the calculations. The palindromes were clustered into five homology groups: C1 (C1+C6+C8+C10+C14+C16), C2 (C2+C11+C15), C3 (C3+C12), C4 (C4+C13), and C5 (C5+C7+C9). The numbers in bold represent the palindrome with highest coverage within each group, which we used as the representative coverage of that homology group.

| **Chimpanzee palindrome** | **Length, bp** | | | | | **Coverage, percentage** | | | |
|---|---|---|---|---|---|---|---|---|---|
| | **Human** | **Bonobo** | **Gorilla** | **Orangutan** | **Chimpanzee*** | **Human** | **Bonobo** | **Gorilla** | **Orangutan** |
| C1 | 36118 | 35368 | 25476 | 37460 | 66916 | **53.98** | **52.85** | **38.07** | **55.98** |
| C2 | 35233 | 111830 | 27957 | 38673 | 141680 | 24.87 | 78.93 | 19.73 | 27.30 |
| C3 | 58885 | 59610 | 57211 | 38076 | 82274 | **71.57** | **72.45** | **69.54** | **46.28** |
| C4 | 119067 | 120124 | 119349 | 116986 | 127172 | **93.63** | **94.46** | **93.85** | **91.99** |
| C5 | 113353 | 107475 | 113915 | 110252 | 136579 | **82.99** | **78.69** | **83.41** | **80.72** |
| C6 | 16015 | 15745 | 6517 | 17378 | 58166 | 27.53 | 27.07 | 11.20 | 29.88 |
| C7 | 45695 | 45809 | 46167 | 45408 | 137290 | 33.28 | 33.37 | 33.63 | 33.07 |
| C8 | 21783 | 21047 | 12228 | 22766 | 64725 | 33.65 | 32.52 | 18.89 | 35.17 |
| C9 | 46890 | 46809 | 47450 | 46382 | 139044 | 33.72 | 33.66 | 34.13 | 33.36 |
| C10 | 16096 | 15957 | 6926 | 17487 | 59322 | 27.13 | 26.90 | 11.68 | 29.48 |
| C11 | 24198 | 105870 | 27840 | 38567 | 123692 | 19.56 | 85.59 | 22.51 | 31.18 |
| C12 | 46539 | 47451 | 46260 | 26584 | 76418 | 60.90 | 62.09 | 60.54 | 34.79 |
| C13 | 117844 | 118277 | 117312 | 116043 | 132034 | 89.25 | 89.58 | 88.85 | 87.89 |
| C14 | 19998 | 19943 | 11045 | 21285 | 47853 | 41.79 | 41.68 | 23.08 | 44.48 |
| C15 | 23050 | 95239 | 25619 | 36280 | 111841 | **20.61** | **85.16**** | **22.91** | **32.44** |
| C16 | 40950 | 33549 | 28037 | 41545 | 76391 | 53.61 | 43.92 | 36.70 | 54.38 |
| C17 | 15364 | 15072 | 12961 | 12978 | 17516 | **87.71** | **86.05** | **74.00** | **74.09** |
| C18 | 6686 | 2582 | 4741 | 6304 | 6888 | **97.07** | **37.49** | **68.83** | **91.52** |
| C19 | 155747 | 156318 | 57117 | 122860 | 168341 | **92.52** | **92.86** | **33.93** | **72.98** |
| Total | 426178 | 488431 | 303092 | 383879 | 637282 | 72.96 | 81.67 | 57.36 | 66.48 |

*Palindrome arm length

**In the case of cluster C2, different palindromes had highest coverage for different species and we considered C15 as a representative because it had the highest coverage for more than one species, while other palindromes had highest coverage for only one species.

Table S7. The copy number for sequences homologous to (A) human and (B) chimpanzee palindromes.

The numbers for bonobo, gorilla, and orangutan were obtained based on median read depth of 1-kb windows homologous to human or chimpanzee palindromes. The copy number for chimpanzee and human were obtained by examining the dotplot of human and chimpanzee Y(1). In brackets, we indicate the known homologs of chimpanzee and human palindromes in human and chimpanzee, respectively (1).

**A**

| Palindrome/ Species | P1 | P2 | P3* | P4 | P5 | P6 | P7 | P8 |
|---|---|---|---|---|---|---|---|---|
| **Human** | 2 | 2 | 2 | 2 | 2 | 2 | 2 | 2 |
| **Chimpanzee** | >2(C3+C4) | >2(C3) | >2(C1 parts) | 0 | >2(C4) | 2(C19) | 2(C18) | 2(C17) |
| **Bonobo** | 1.64 | 1.46 | 1.62 | 0 | 2.13 | 1.42 | 0.70 | 1.98 |
| **Gorilla** | 1.99 | 1.99 | 1.69 | 2.49 | 1.91 | 1.4 | 1.23 | 0.88 |
| **Orangutan** | 6.52 | 13.13 | 1.05 | 5.67 | 7.29 | 1.06 | 1.04 | 1.80 |

*Note: We assume that some parts of human palindrome P3 in chimpanzee are multi-copy (those that share homology with C1), while others are single-copy.

**B**

| Palindrome/ Species | C1 group | C2 group | C3 group | C4 group | C5 group | C17 | C18 | C19 |
|---|---|---|---|---|---|---|---|---|
| **Human** | 2(P3 parts) | 0 | >2(P1,P2) | >2(P5,P1) | 1 | 2 (P8) | 2(P7) | 2(P6) |
| **Chimpanzee** | >2 | >2 | >2 | >2 | >2 | 2 | 2 | 2 |
| **Bonobo** | 13.29 | 8.42 | 1.18 | 1.41 | 9.31 | 2.27 | 0.80 | 1.25 |
| **Gorilla** | 1.87 | 2.54 | 2.01 | 2.04 | 1.12 | 0.93 | 1.27 | 1.26 |
| **Orangutan** | 1.02 | 6.07 | 12.13 | 7.85 | 1.07 | 1.87 | 1.02 | 1.02 |

#### Table S8. Summary of Y chromosome species-specific sequences.

The copy number of species-specific blocks in multi-species alignments. Only >100-bp blocks were analyzed to compute the percentages (shown in parentheses) from the total species-specific sequence. CN - copy number.

| Species | Total length | In blocks with CN≤1.33 (i.e. CN≈1) | In blocks with 1.33<CN≤1.66 (CN between 1 and 2, uncertain) | In blocks with 1.66<CN≤2.5 (i.e. CN≈2) | In blocks with CN>2.5 (i.e. CN>2) |
|---|---|---|---|---|---|
| **Bonobo** | 9,473,169 bp | 1,180,899 bp (12.47%) | 630,172 bp (6.65%) | 1,673,854 bp (17.67%) | 5,976,284 bp (63.09%) |
| **Gorilla** | 1,632,223 bp | 704,943 bp (43.19%) | 202,842 bp (12.43%) | 218,085 bp (13.36%) | 496,387 bp (30.41%) |
| **Orangutan** | 3,520,105 bp | 2,247,989 bp (63.46%) | 136,390 bp (3.85%) | 190,607 bp (5.41%) | 945,119 bp (26.68%) |

Table S9. The number of observed chromatin contacts, followed by the chromatin interactions weighted by their probability (multi-mapping reads are allocated probabilistically (46)). The density of chromatin interactions is higher in palindromes. The table is based on human iPSC data (47). See also Fig. S8.

| HUMAN | total number of chromatin interactions | weighted number of chromatin interactions | region length [Mb] | **density of weighted interactions [per Mb]** |
|---|---|---|---|---|
| palindrome interactions | 82,619 | 14,362 | 6 | **2,489** |
| mixed interactions | 70,533 | 12,111 | - | - |
| other interactions | 181,178 | 26,064 | 24 | **1,086** |
| sum | 334,330 | 52,536 | 30 | |

Table S10. The generated-here and previously unpublished sequencing datasets used for assembly generation and classification, including their summary information.

| Species | Read type | Source | Individual ID | Insert size (bp) | Read count | Read length (bp) | Sequencing technology |
|---|---|---|---|---|---|---|---|
| Bonobo | paired-end | Whole-genome | PR00251 | 1,000 | 319,637,420 | 251 | Illumina HiSeq2500 |
| Bonobo | mate-pair | Whole-genome | PR00251 | 8,000 | 120,181,539 | 100 | Illumina HiSeq2500 |
| Bonobo | long reads | Y flow-sorted | Ppa_MFS | >7,000 | 818,697 | variable | PacBio |
| Orangutan | paired-end | Whole-genome | 1991-0051 | 1,000 | 303,928,291 | 251 | Illumina HiSeq2500 |
| Orangutan | synthetic long reads | Whole-genome | AG06213 | ~350 bp | 437,361,894 | 151 | 10X Genomics |
| Orangutan | mate-pair | Whole-genome | 1991-0051 | 8,000 | 120,384,643 | 151 | Illumina HiSeq2500 |
| Gorilla | paired-end | Whole-genome | KB3781 | 550 bp | 120,740,133 | 150 | NextSeq |

*Before any trimming, adapter removal or quality filtering. For paired reads, the number of read pairs is listed.

#### Table S11. The number of variants before and after the polishing step.

The haploid variants reported by FREEBAYES (14) were filtered to retain only high quality calls (QUAL≥ 30). The polishing step reduces the number of variants present in the assembly.

| ORANGUTAN | Before the polishing | After the polishing |
|---|---|---|
| All called variants | 294,544 | 286,782 |
| Called variants with QUAL≥30 | 38,597 | 4,537 |
| BONOBO | Before the polishing | After the polishing |
| All called variants | 414,617 | 408,580 |
| Called variants with QUAL≥30 | 52,158 | 5,318 |

### Table S12. The coordinates of palindromes on panTro6 and hg38 Y chromosome.

The coordinates were obtained from (4) and updated to the current version of chimpanzee Y, panTro6.

**A**

| Palindrome | Start | End | The end of left arm (approximated) |
|---|---|---|---|
| P1 | 23359067 | 26311550 | 24822577 |
| P2 | 23061889 | 23358813 | 23208197 |
| P3 | 21924954 | 22661453 | 22208730 |
| P4 | 18450291 | 18870104 | 18640356 |
| P5 | 17455877 | 18450126 | 17951255 |
| P6 | 16159590 | 16425757 | 16269541 |
| P7 | 15874906 | 15904894 | 15883575 |
| P8 | 13984498 | 14058230 | 14019652 |

**B**

| Chimpanzee palindrome | Start | End | Amplicon color following (1) | Approximate arm length (half of palindrome) | The end of left arm (approximated) |
|---|---|---|---|---|---|
| **C1** | 1759451 | 2,053,069 | Pink | 146809 | 1906260 |
| **C2** | 2298081 | 2,984,818 | Blue | 343368.5 | 2641450 |
| **C3** | 3587737 | 3,925,944 | Red | 169103.5 | 3756841 |
| **C4** | 4669973 | 5,444,881 | Gold | 387454 | 5057427 |
| **C5** | 8651546 | 9,099,775 | Turquoise | 224114.5 | 8875661 |
| **C6** | 9099776 | 9,383,863 | Pink | 142043.5 | 9241820 |
| **C7** | 9383864 | 9,832,093 | Turquoise | 224114.5 | 9607979 |
| **C8** | 9832094 | 10,116,181 | Pink | 142043.5 | 9974138 |
| **C9** | 10116182 | 10,564,411 | Turquoise | 224114.5 | 10340297 |
| **C10** | 10564412 | 10,851,248 | Pink | 143418 | 10707830 |
| **C11** | 11060051 | 11,674,261 | Blue | 307105 | 11367156 |
| **C12** | 12651797 | 12,963,129 | Red | 155666 | 12807463 |
| **C13** | 13340000 | 14,084,660 | Gold | 372330 | 13712330 |
| **C14** | 14798116 | 15,024,316 | Pink | 113100 | 14911216 |
| **C15** | 15493746 | 16,056,815 | Blue | 281534.5 | 15775281 |
| **C16** | 16477310 | 16,791,733 | Pink | 157211.5 | 16634522 |
| **C17** | 21591500 | 21,671,300 | | 39900 | 21631400 |
| **C18** | 23517939 | 23,546,819 | | 14440 | 23532379 |
| **C19** | 23807052 | 24,577,396 | | 385172 | 24192224 |

Table S13. The list of ENCODE datasets analyzed in the search for regulatory elements in P6 and P7.

| Experiment | Unfiltered BAM | Target | Tissue | Link |
|---|---|---|---|---|
| ENCSR000ALB | ENCFF735TGN | H3K27Ac | HUVEC | https://www.encodeproject.org/experiments/ENCSR000ALB/ |
| ENCSR000AKL | ENCFF322MOQ | H3K4me1 | HUVEC | https://www.encodeproject.org/experiments/ENCSR000AKL/ |
| ENCSR000ALG | ENCFF261CBZ | Control | HUVEC | https://www.encodeproject.org/experiments/ENCSR000ALG/ |
| ENCSR000EOQ | ENCFF042VZB | Dnase-seq | HUVEC | https://www.encodeproject.org/experiments/ENCSR000EOQ/ |
| ENCSR112ALD | ENCFF319GEZ | CREB1 | HepG2 | https://www.encodeproject.org/experiments/ENCSR112ALD/ |
| ENCSR136ZQZ | ENCFF807LQS | H3K27Ac | Testis | https://www.encodeproject.org/experiments/ENCSR136ZQZ/ |
| ENCSR956VQB | ENCFF077NRU | H3k4me1 | Testis | https://www.encodeproject.org/experiments/ENCSR956VQB/ |
| ENCSR215WNN | ENCFF511LQO | Control | Testis | https://www.encodeproject.org/experiments/ENCSR215WNN/ |
| ENCSR729DRB | ENCFF639PHQ | Dnase-seq | Testis | https://www.encodeproject.org/experiments/ENCSR729DRB/ |

### Supplemental Figures

Figure S1. Flowcharts for the assemblies of (A) bonobo, (B) Sumatran orangutan, and (C) gorilla.

Blue: input datasets, green: software tools, pink: processed datasets.

**A. Bonobo**

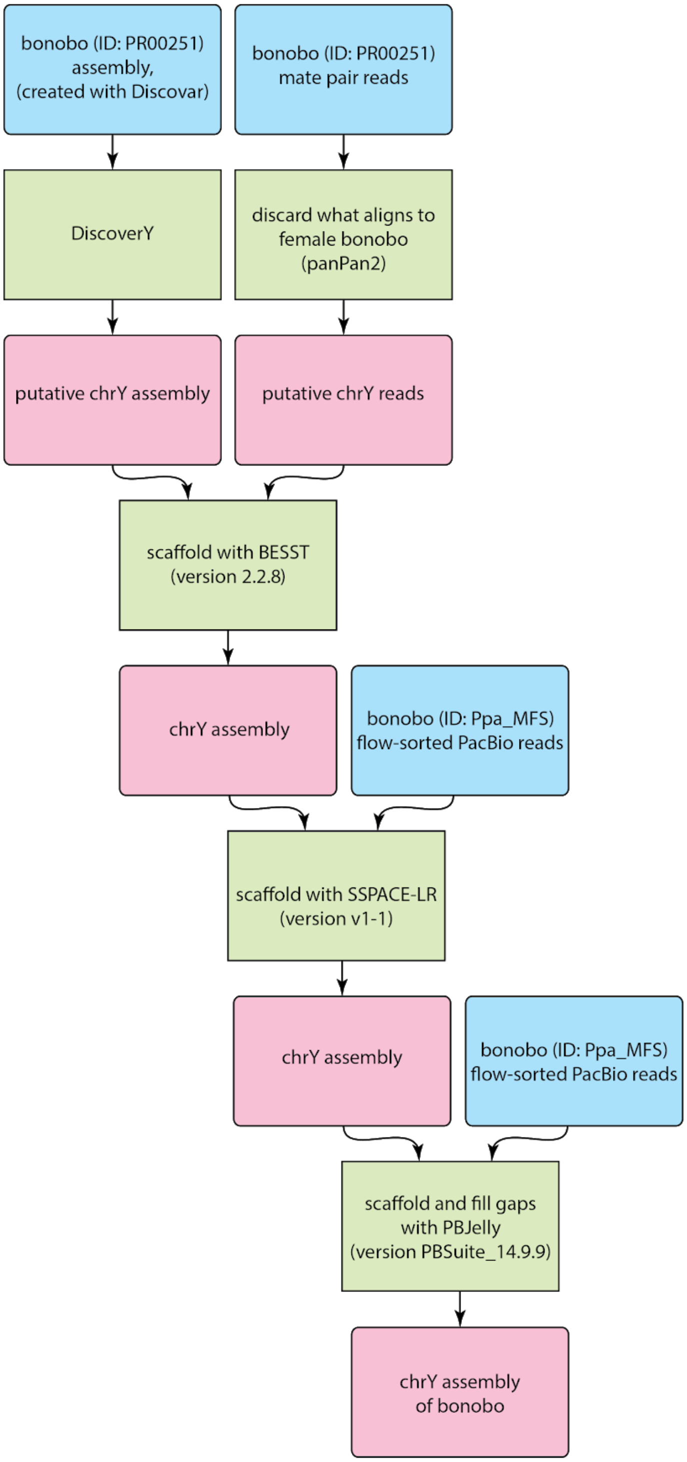

**B. Sumatran orangutan**

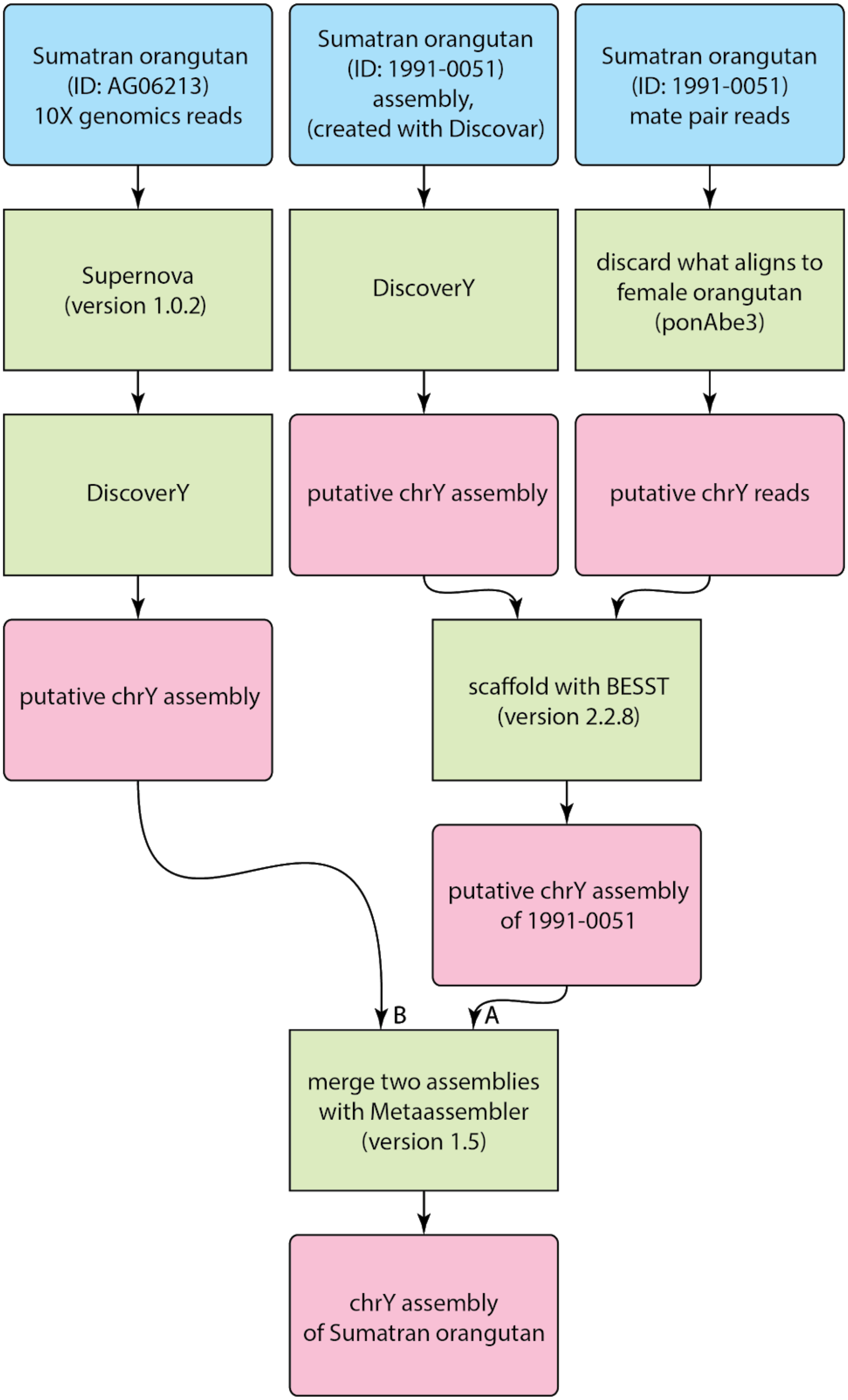

**C. Gorilla**

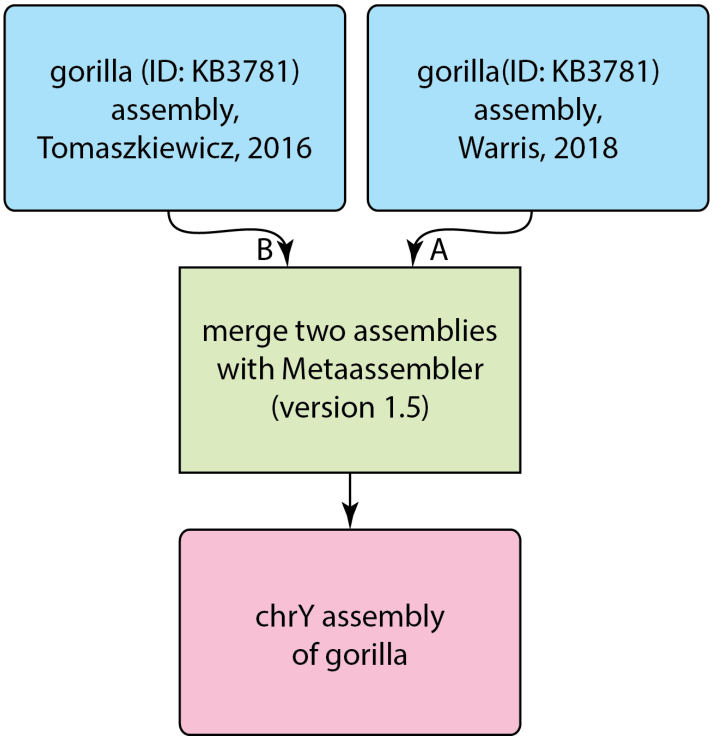

gorilla (ID: KB3781) assembly, Tomaszkiewicz, 2016
gorilla(ID: KB3781) assembly, Warris, 2018
B
A
merge two assemblies with Metaassembler (version 1.5)
chrY assembly of gorilla

Figure S2A. Sequence comparisons between human (hg38) and bonobo, gorilla and orangutan Y chromosome assemblies.

The order of scaffolds and/or contigs was modified in each assembly to match the human Y chromosome using MAUVE (65) v.2015-02-25. The heterochromatic portion of the *q* arm of the human Y chromosome was omitted. Dot plots were generated with GEPARD v.1.40 (36) using word length 50.

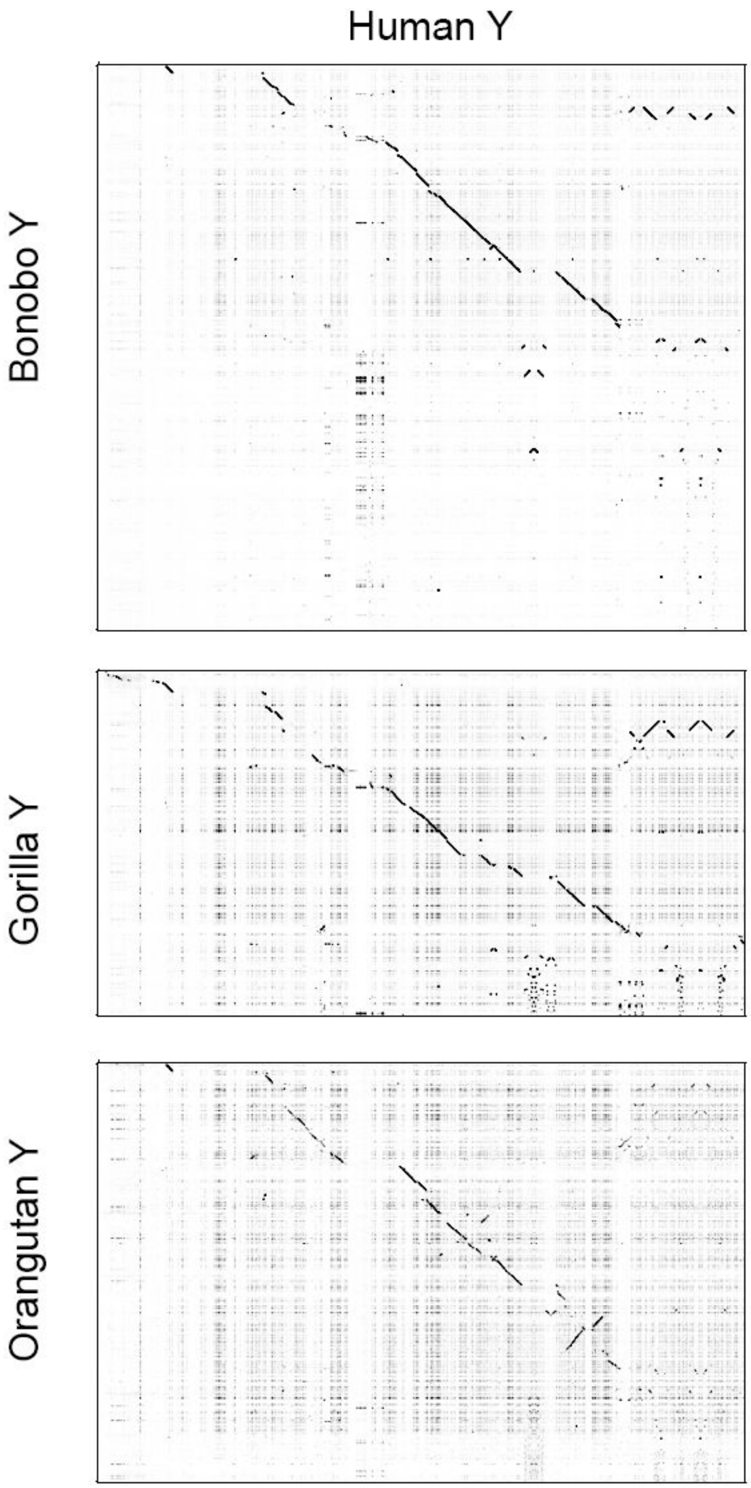

#### Figure S2B. Sequence comparisons between chimpanzee (panTro6) and bonobo, gorilla and Sumatran orangutan Y chromosome assemblies.

The order of scaffolds and/or contigs was modified in each assembly to match the chimpanzee Y chromosome using MAUVE (65) v.2015-02-25. Dot plot was generated with GEPARD version 1.40 (36) using word length 50.

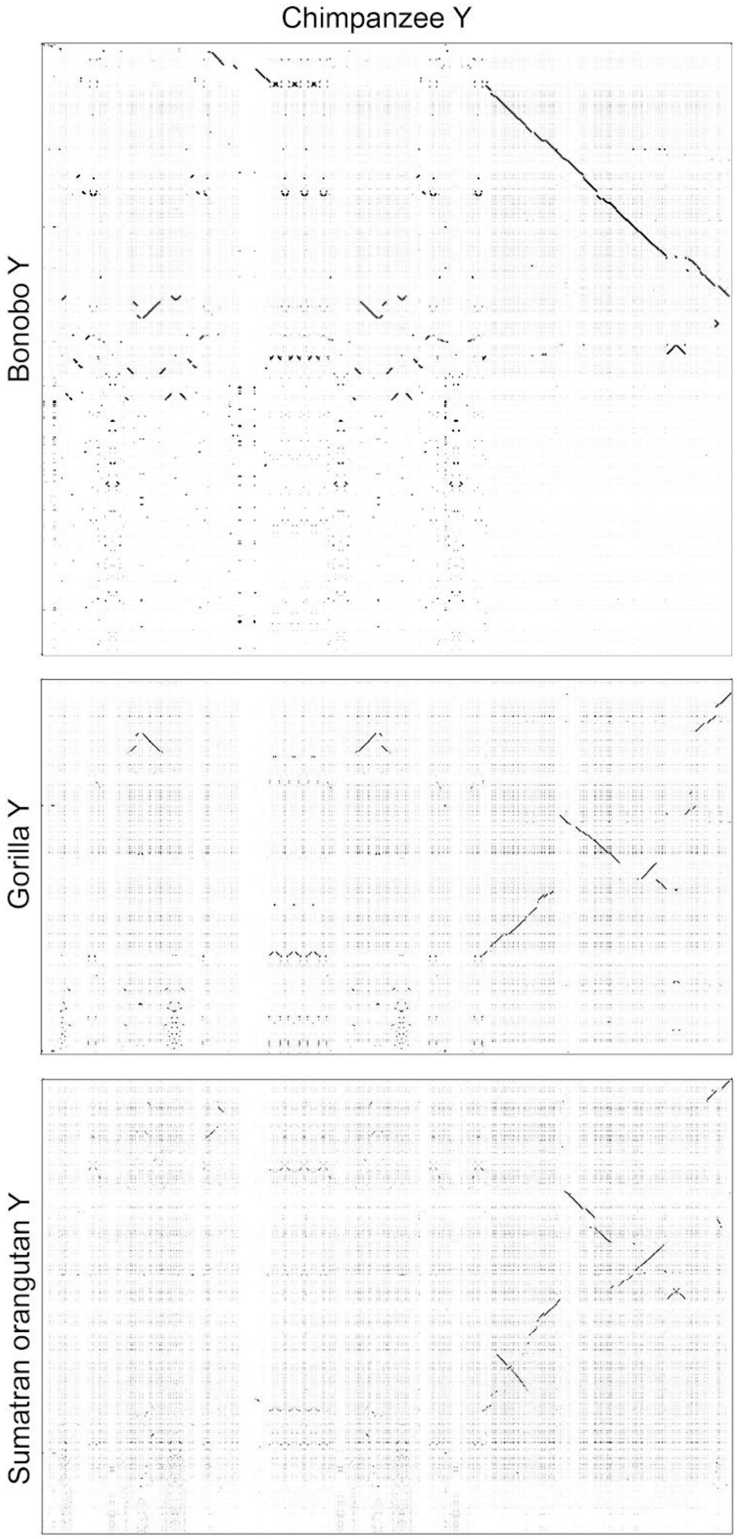

### Figure S3. Protein-coding gene sequence retrieval in the new Y assemblies.

**(A)** For evaluation purposes, we aligned the scaffolds from each of bonobo, Sumatran orangutan and gorilla Y chromosome assemblies to species-specific or closest-species-specific reference coding sequences using BWA-MEM (v.0.7.10) (10). Next, we visualized the alignment results in Integrative Genomics Viewer (IGV) (v.2.3.72). Consensus sequences were retrieved to evaluate sequence coverage over the reference sequence (in percentage) using BLAST (31). The results were represented as heatmaps using pheatmap package in R. **(B)** Table of accession numbers used as queries.

**A**

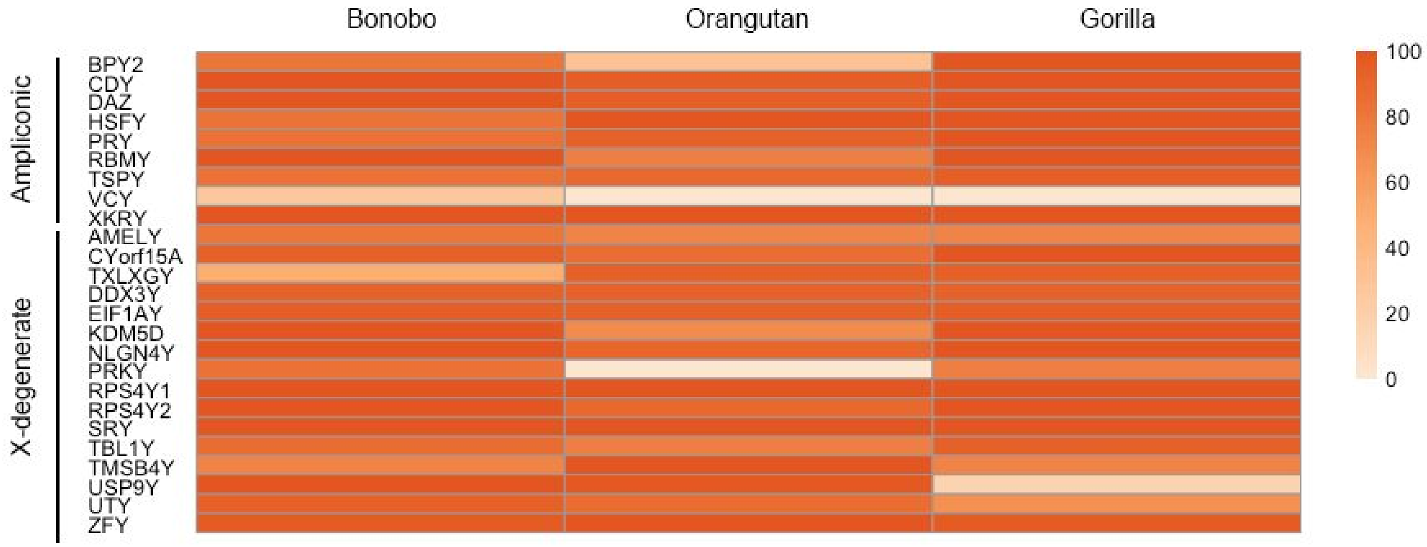

**B**

| Coding sequence | Bonobo | Orangutan | Gorilla |
|---|---|---|---|
| ***BPY2*** | AY958084.1 | KP141770.1 | GATR01000016.1 |
| ***CDY*** | AY958081.1 | KP141772.1 | GATR01000022.1 |
| ***DAZ*** | AY958083.1 | KP141773.1 | GATR01000021.1 |
| ***HSFY*** | CCDS35475.1 | KP141774.1 | GATR01000004.1 |
| ***PRY*** | CCDS14799.1 | KP141776.1 | GATR01000017.1 |
| ***RBMY*** | AH014838.2 | KP141777.1 | GATR01000007.1 |
| ***TSPY*** | AY958082.1 | KP141780.1 | GATR01000012.1 |
| ***VCY*** | XM_003318999.5 | CCDS56617.1 | CCDS56617.1 |
| ***XKRY*** | NM_004677.2 | NM_004677.2 | NM_004677.2 |
| ***AMELY*** | NM_001102459.1 | ENST00000215479.10 | FJ532255.1 |
| ***CYorf15A*** | AY633113.1 | NR_045128.1 | FJ532256.1 |
| ***TXLNGY*** | NR_045129.1 | GATK01000021.1 | FJ532257.1 |

| | | | |
|---|---|---|---|
| ***DDX3Y*** | AY633112.1 | NM_001131248.1 | FJ532258.1 |
| ***EIF1AY*** | AY633115.1 | GATK01000002.1 | FJ532259 |
| ***KDM5D*** | AY736376.1 | GATK01000003.1 | FJ532260.1 |
| ***NLGN4Y*** | AY728015.1 | KP141775.1 | FJ532261 |
| ***PRKY*** | AY728014.1 | ENST00000533551.5 | FJ532262 |
| ***RPS4Y1*** | AY633110.1 | GATK01000004.1 | FJ532263.1 |
| ***RPS4Y2*** | AY633111.1 | KP141778.1 | FJ532264 |
| ***SRY*** | AY679780.1 | KP141779.1 | X86382.1 |
| ***TBL1Y*** | ENST00000383032.6 | GATK01000018.1 | FJ532265 |
| ***TMSB4Y*** | ENST00000284856.4 | GATK01000007.1 | FJ532266 |
| ***USP9Y*** | ENST00000338981.7 | GATK01000005.1 | FJ532267 |
| ***UTY*** | AY679781.1 | GATK01000020.1 | FJ532268.1 |
| ***ZFY*** | AY679779.1 | GATK01000006.1 | AY913764 |

Figure S4. (A, C, E) Thresholds used for classification of windows into X-degenerate versus ampliconic, and (B, D, F) average copy number for overlapping 5-kb windows. X-degenerate windows are shown in yellow, whereas ampliconic windows are shown in blue (or turquoise

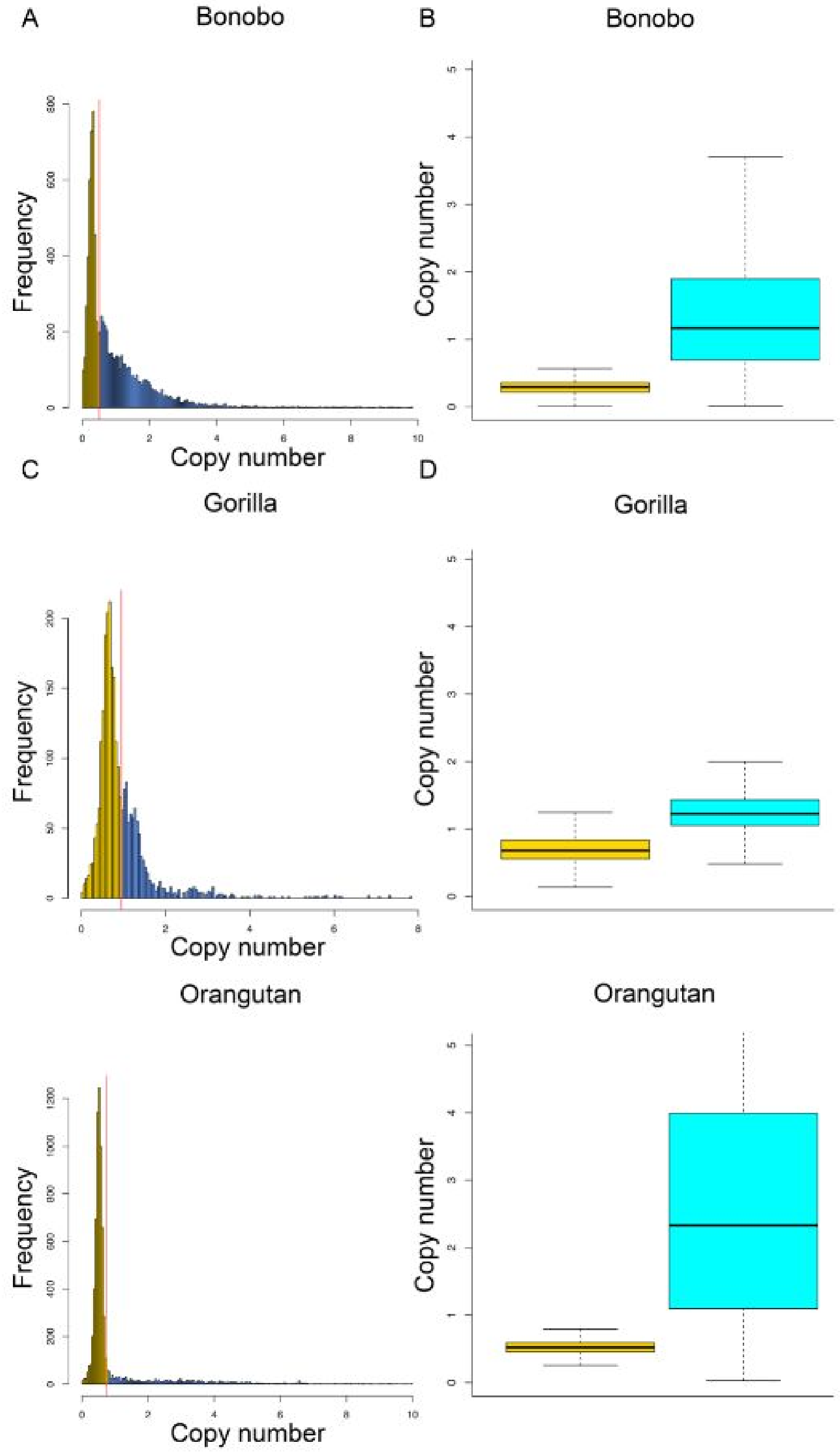

Figure S5. Shared and lineage-specific sequences in multi-species alignments.
Counts of aligned bases in each set of species. For example, the first five bars reflect alignments involving human, chimpanzee, bonobo, gorilla, and orangutan; the sixth through ninth bars reflect alignments involving human, chimpanzee, bonobo, and gorilla but not orangutan, etc.; the last five columns reflect species-specific sequences.

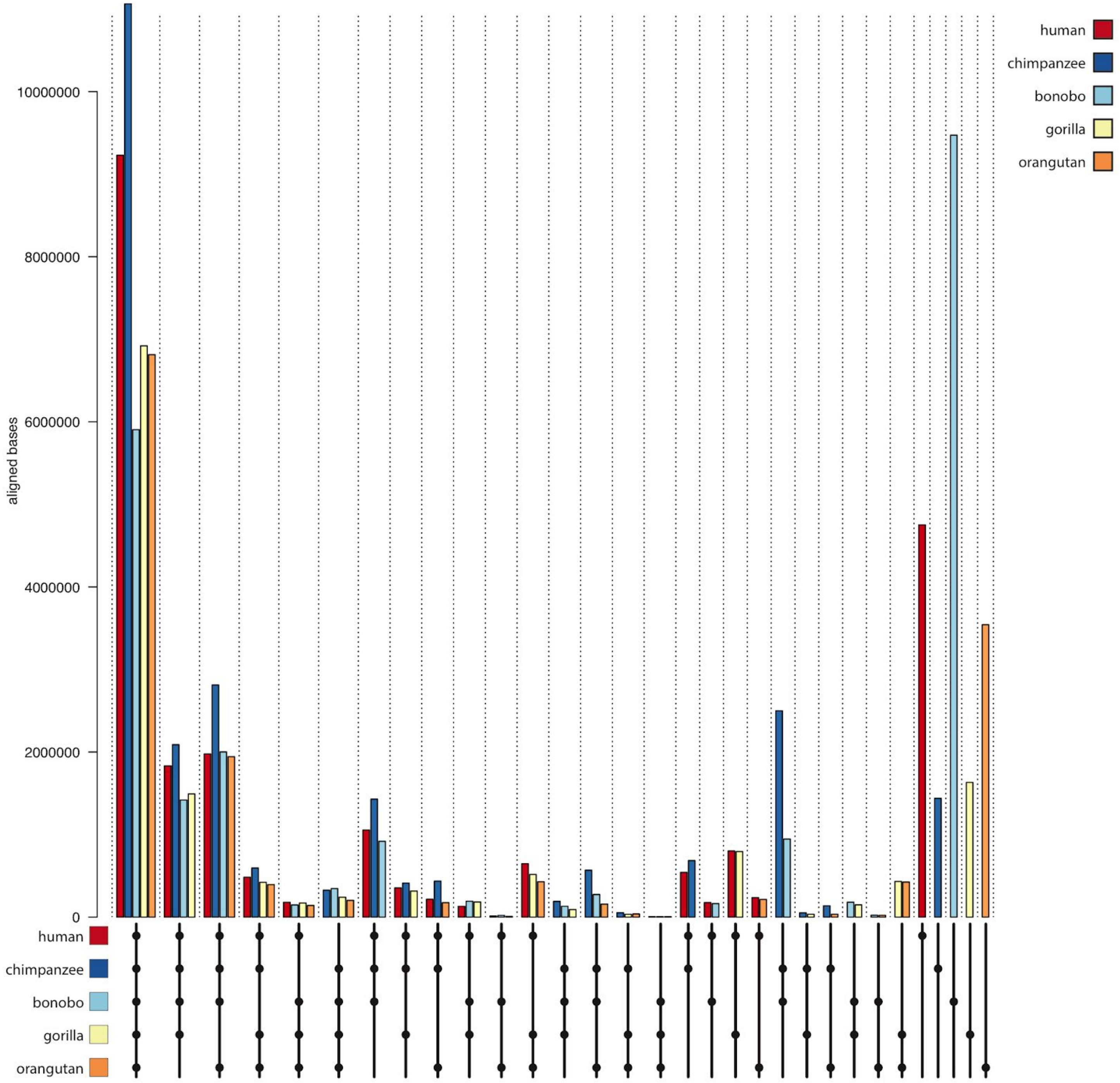

### Figure S6. Reconstructed gene content of great apes.

The first six rows have information about the gene content of great apes and macaque, which were used as an input for the model of Iwasaki and Takagi (33). The other rows were reconstructed by the model. BC - common ancestor of bonobo and chimpanzee; BCH - common ancestor of bonobo, chimpanzee, and human; BCHG - common ancestor of bonobo, chimpanzee, human, and gorillas. GA - common ancestor of great apes. We defined the presence of *RPS4Y2* and *MXRA5Y* in bonobo and orangutan based on AUGUSTUS, and Y chromosome specific testis transcriptome assembly results (29). The presence of *RPS4Y2* gene was confirmed in bonobo through gene prediction (shares 100% identity with chimpanzee *RPS4Y2*) and assembled transcript sequences (shares 99.6% identity with chimpanzee *RPS4Y2*). *MXRA5Y*, which is psedogenized in human and chimpanzee (66), was missing in orangutan (no gene prediction or transcript found) and pseudogenized in bonobo (gene prediction annotated as X chromosome homolog *MXRA5* and missing the first three exons of *MXRA5* in its sequence). In the case of gorilla, we did not find its transcript in transcriptome assembly and a BLAT search of human *MXRA5Y* gene (NC_000024.10:11952465-11993293) resulted in a 12-kb long hit which is around 20% of the gene. We assumed the gene is lost in gorilla as well.

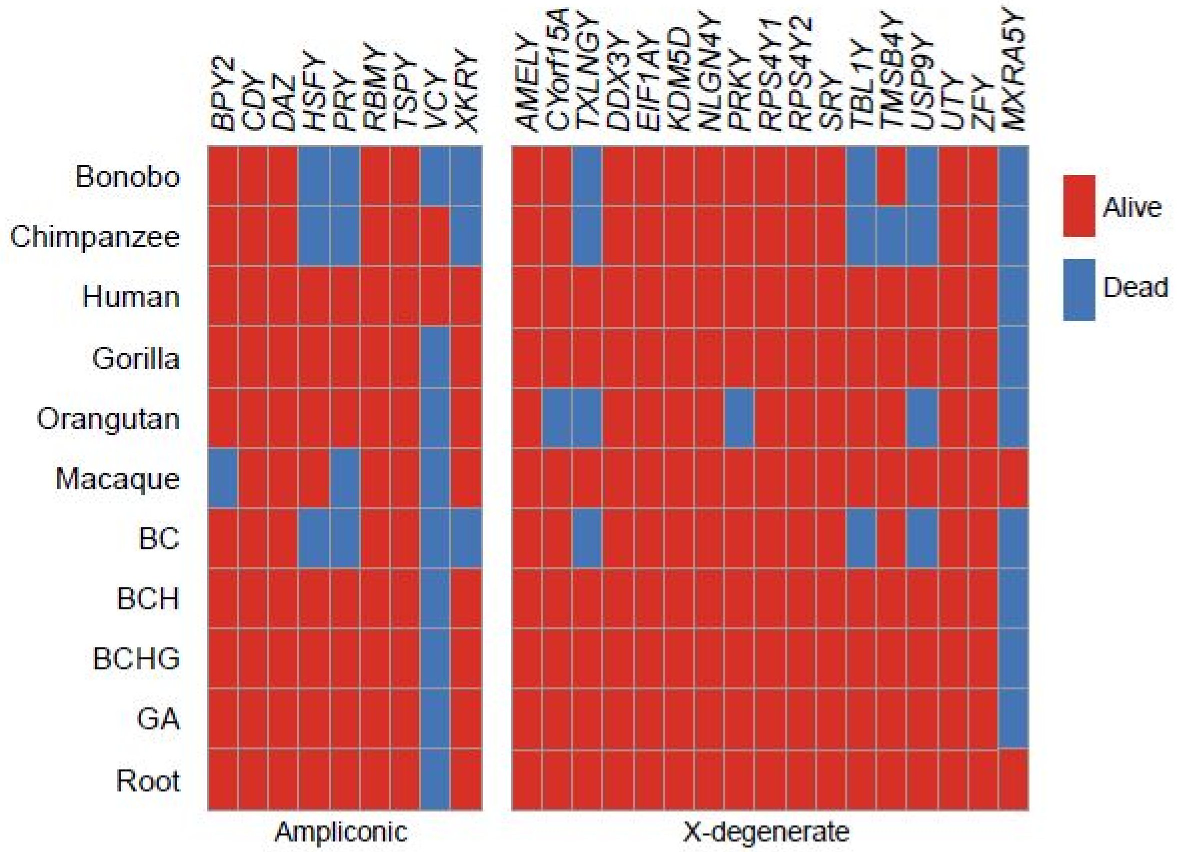

Figure S7. IGV screen shots of peaks of DNase-seq, H3K4me1 and H3K27ac marks on human palindrome P6, and of CREB1 on human palindrome P7.

**A**. Peaks on both arms of P6 are shown within the blue and grey boxes. **B**. Zoom-in view of peaks on the left arm of palindrome P6. The coverage track represents the depth of coverage and peaks track represents the peaks identified by MACS2 (44). **C**. Zoom-in view of peaks on the right arm of palindrome P6. The coverage track (top) represents the depth of coverage, and the peaks track (bottom) represents the peaks identified by MACS2 (44). **D.** Peaks on both arms of palindrome P7 are shown. The coverage track (top) represents the depth of coverage and the peaks track (bottom) represents the peaks identified by MACS2 (44). See Table S12 for the coordinates of P7.

A

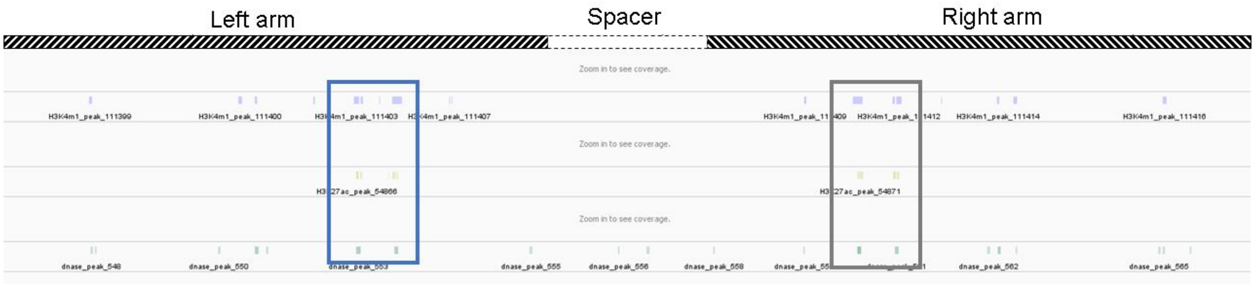

B

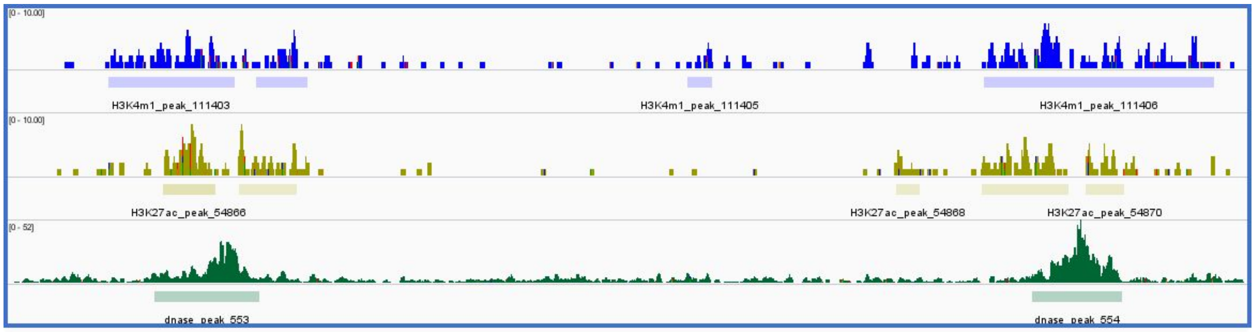

C

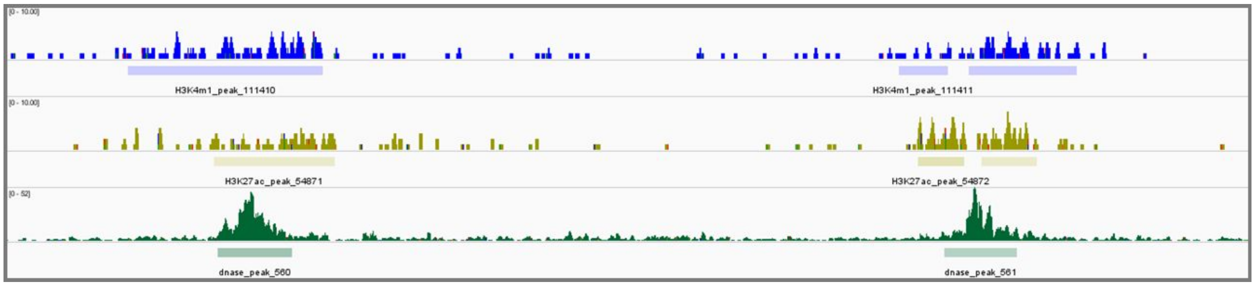

D

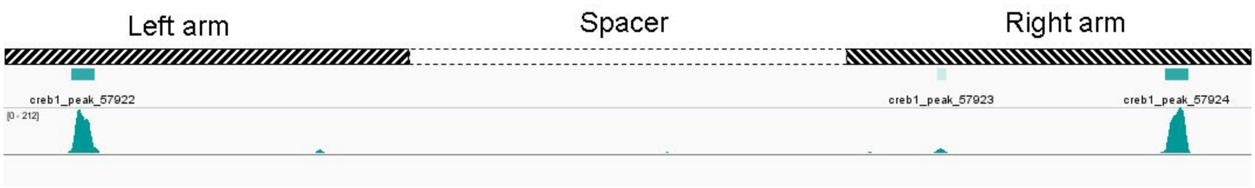

#### Figure S8. Chromatin interactions on the human Y chromosome.

**A**. Chromatin contacts split by groups: palindrome-palindrome contacts, palindrome-other (i.e. mixed) and other-other (chromatin interactions in which palindromes are not involved). **B**. The probability of interactions as estimated by (46); the probability is the highest for the palindromic group, in which both reads from a pair fall into human palindromes. The table is based on human iPSC data (47). See also Table S9.

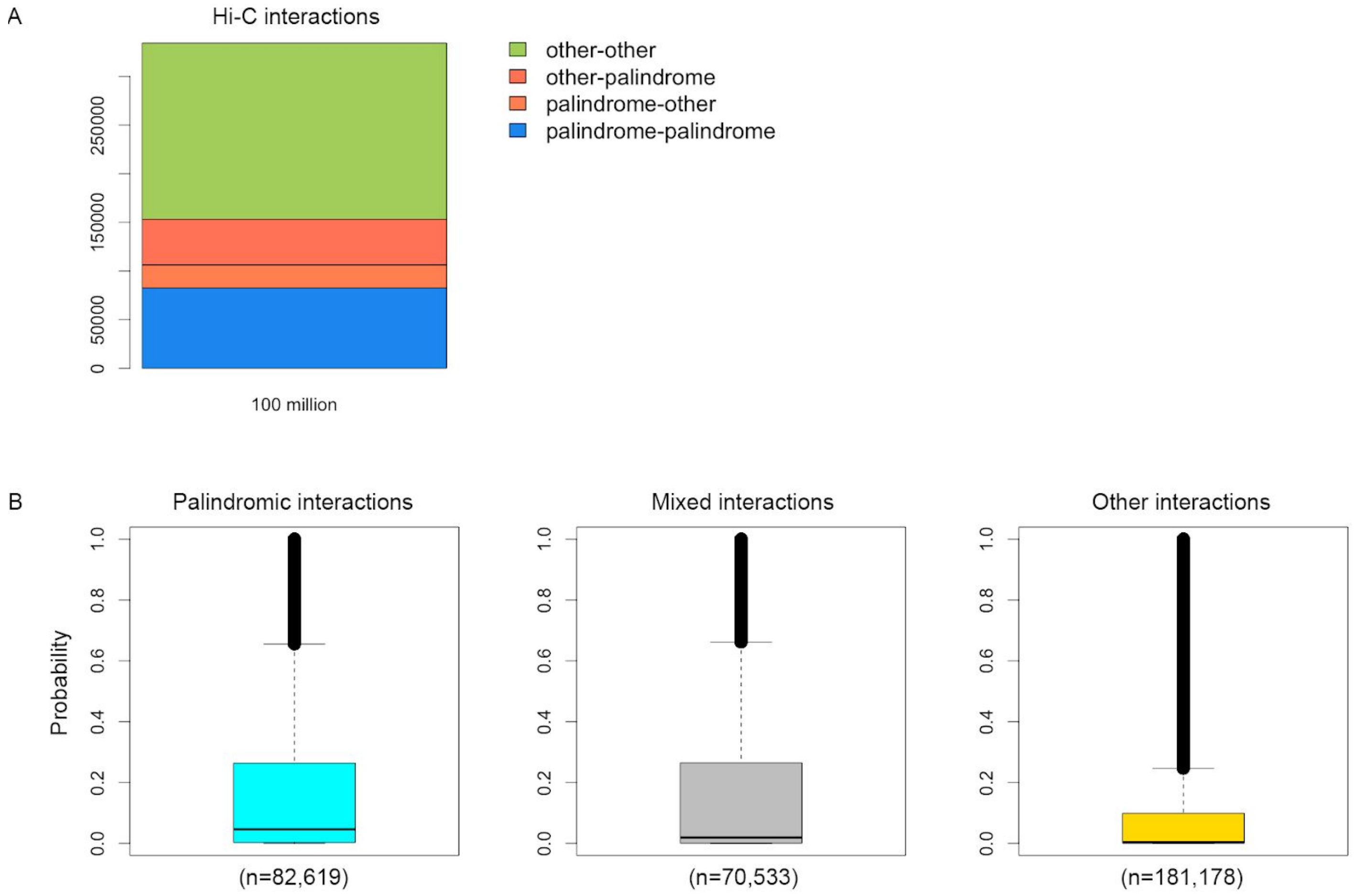

Figure S9. Hi-C contact map generated for Human Umbilical Vein Cells (HUVEC), using the data from (49).
See color coding explained in Fig. 4.

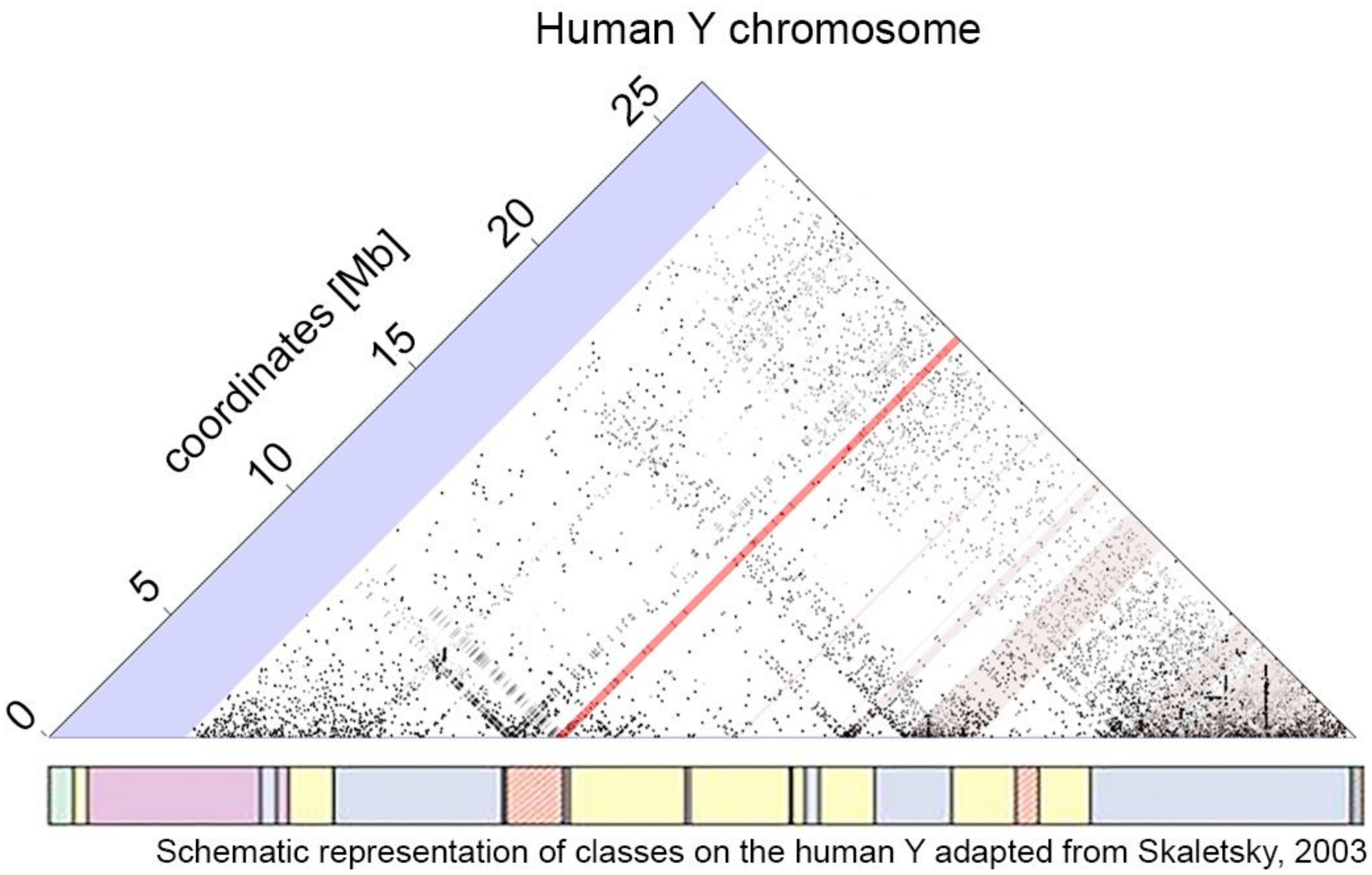
